# Modeling the pathological brain-gut axis in Parkinson’s disease using human iPSC derived brain-intestinal assembloids

**DOI:** 10.64898/2026.02.07.704302

**Authors:** Akihiro Yamagchi, Kei-ichi Ishikawa, Wado Akamatsu

## Abstract

Parkinson’ disease is increasingly recognized as a systemic disorder involving pathological communication along the brain–gut axis, yet the cellular mechanisms underlying inter-organ α-synuclein propagation remain poorly defined. Here, we establish a human induced pluripotent stem cell–derived brain–intestinal assembloid that integrates a developmentally specified medullary hindbrain organoid with an enteric neuron–enriched intestinal organoid. Using this model, we show the propagation of α-synuclein from brainstem neurons to enteric neurons. Ligand–receptor analysis combined with genetic perturbation identifies NRXN1–NLGN2 signaling as a key mediator of brain-to-gut α-synuclein propagation, with NRXN1 being upregulated in Parkinson’s disease patient datasets *in vivo*. Together, these findings establish a human stem cell–based platform for investigating brain–gut communication and uncover molecular mechanisms underlying α-synuclein propagation in Parkinson’s disease.

## INTRODUCTION

Parkinson’s disease (PD) is a neurodegenerative disorder characterized by the progressive accumulation of misfolded α-synuclein (α-syn) aggregates, forming Lewy bodies.^1^ While motor symptoms such as akinesia and rigidity define the clinical diagnosis, non-motor symptoms, including constipation, often emerge decades before the onset of motor manifestations.^2^ This temporal dissociation has increasingly drawn attention to the gastrointestinal tract as a potential initiating site of PD pathogenesis.^1,3^

Previous studies suggest that pathogenic α-syn can propagate across anatomically connected neuronal networks.^4,5^ In this context, the vagus nerve, which directly links the enteric nervous system (ENS) to the dorsal motor nucleus of the vagus (DMX), is thought to serve as a primary conduit for this propagation.^6,7^ Gut-derived α-syn has been shown to ascend to the brainstem in rodent models, supporting the Braak hypothesis that PD pathology may originate peripherally before invading central circuits.^8,9^ Notably, epidemiological studies reporting reduced PD risk following truncal vagotomy further implicate the vagal pathway as a causal route for disease propagation.^10,11^ Conversely, descending propagation of α-syn from the central nervous system to peripheral autonomic structures has also been demonstrated,^12,13^ underscoring the bidirectional nature of brain–gut communication and its contribution to the systemic features of PD. Despite these advances, the cellular and molecular mechanisms that enable α-syn to traverse organ boundaries remain largely undefined.

Several methodological limitations hinder progress. Rodent models dominate current research on the brain–gut axis, yet substantial interspecies differences in microbiome composition, immune physiology, and brainstem organization restrict their translational relevance.^14,15^ Human post-mortem studies have identified α-syn pathology in both intestinal and brainstem tissues, but these static observations cannot capture the early, dynamic processes underlying pathological initiation and spread. More recently, microfluidic “organ-on-a-chip” systems have begun to reconstruct bidirectional gut–brain communication *in vitro*,^16^ but the two-dimensional (2D) architecture lacks the regional patterning and cytoarchitectural complexity required to model vagal circuitry.

Human-induced pluripotent stem cell (iPSC)–derived organoid and assembloid technologies now enable the reconstruction of multicellular circuits with developmental and regional fidelity.^17,18^ Recent studies have shown that brain–brain assembloids can recapitulate α-syn propagation,^19,20^ and that gut–visceral sensory ganglion assembloids can model amyloid-β transmission along the gut–brain axis.^21^ However, no existing human model faithfully reconstructs the vagal–enteric neuroanatomy, nor has any system successfully reproduced α-syn propagation along the vagus nerve. A physiologically structured model that integrates region-specific brainstem neurons with an innervated intestinal microenvironment is therefore critically needed to dissect mechanisms of inter-organ α-syn transmission.

Here, we developed a human iPSC-derived brain–intestinal assembloid that reconstructs medullary hindbrain circuitry and an innervated intestinal compartment within an integrated 3D framework. Using this model, we recapitulated the descending propagation of pathological α-syn from the brainstem to the gut. By combining single-cell transcriptomics, computational ligand–receptor analysis, and genetic perturbation, we identify a previously unrecognized signaling axis that mediates inter-organ α-syn transfer. Collectively, these findings provide a mechanistic foundation for understanding human brain–gut α-syn propagation and highlight potential therapeutic targets for early intervention in PD.

## RESULTS

### Generation of Medullary Hindbrain Organoids with DMX Identity

To model the vagal origin of PD-associated pathology, we first optimized a differentiation protocol to generate medullary hindbrain organoids (mHBOs). As WNT and retinoic acid (RA) signaling act as key caudalization cues during hindbrain development,^22,23^ we systematically optimized the concentrations of CHIR99021 and RA to induce a posterior hindbrain identity capable of producing DMX-like visceral motor neurons (Figure 1A). Among the conditions tested, a defined CHIR–RA window yielded the most robust induction of medullary markers, including HOXB4 and PHOX2B, as assessed by RT–qPCR at day 11 (Figure 1B). These results were reproducible across independent differentiation batches and multiple iPSC lines (Figure S1). Consistent with transcriptional profiles, immunostaining confirmed protein-level expression of GBX2, HOXB4, and PHOX2B, while showing minimal expression of more rostral (OTX2) or caudal (HOXC6) determinants, validating accurate regional specification (Figure 1C).

**Figure 1.**
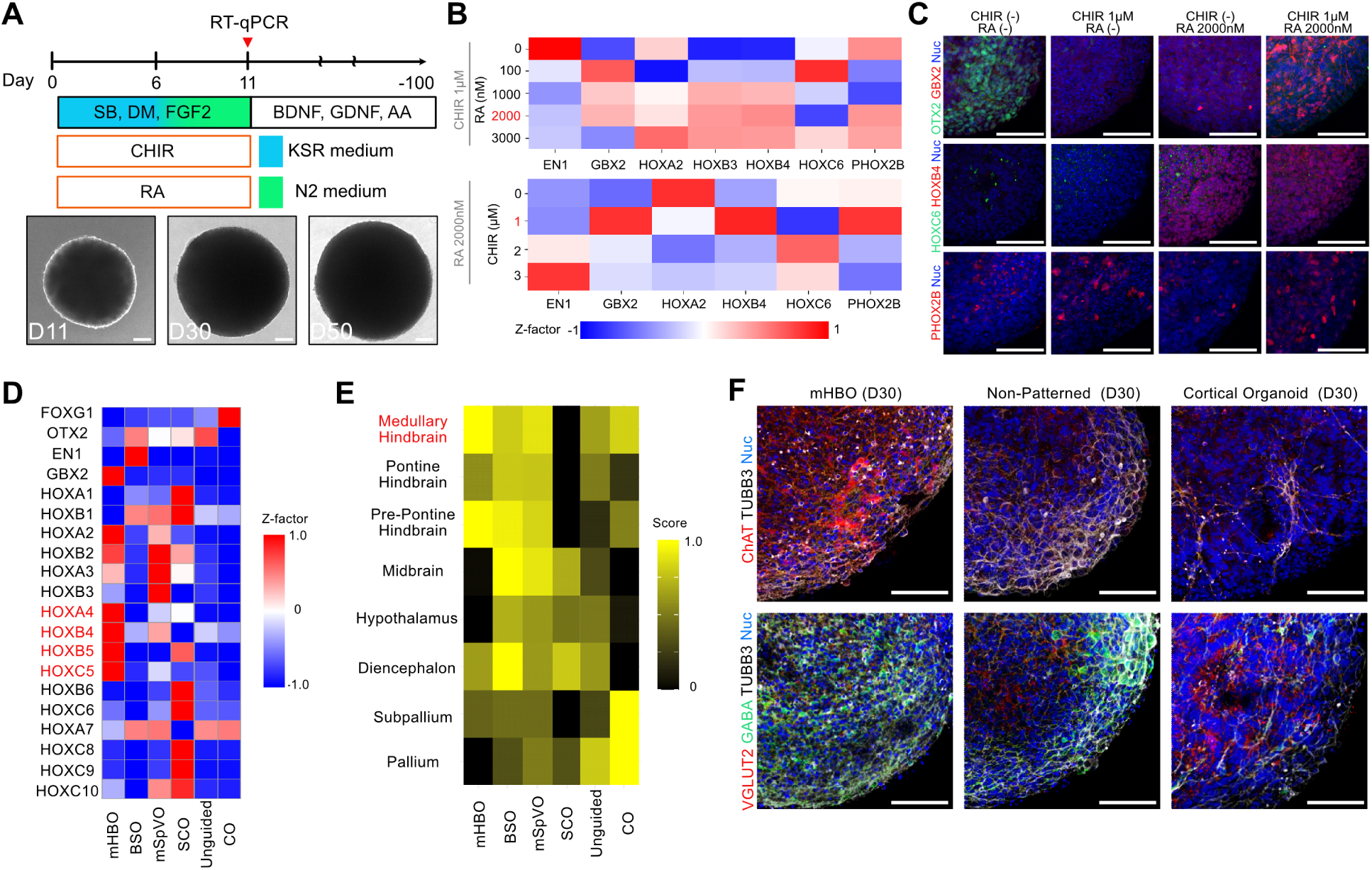
Morphogen optimization for hiPSC-derived mHBO generation. (A) Schematic of the directed differentiation protocol used to generate medullary hindbrain organoids (mHBOs) (top). Representative brightfield images show organoid morphology across differentiation time points (bottom). For morphogen optimization, CHIR99021 (CHIR) and retinoic acid (RA) concentrations were titrated during the patterning window and marker expression was assayed at day 11. SB, SB431542; DM, dorsomorphin; BDNF, brain-derived neurotrophic factor; GDNF, glial cell line-derived neurotrophic factor; AA, ascorbic acid. Scale bar, 100 μm. (B) Heatmap of scaled RT–qPCR marker expression (day 11) from optimized conditions (z-scaled per gene). Conditions that robustly induced medullary markers are highlighted. See also Figure S1. (C) Immunofluorescence staining for OTX2, GBX2, HOXB4, HOXC6, and PHOX2B in day 11 organoids. Scale bars, 100 μm. (D) Heatmap showing the expression of region-specific genes in mHBOs compared with published datasets based on bulk RNA-seq. BSO, brain stem organoid; mSpVO, medullary spinal trigeminal nucleus-like organoid; SCO, spinal cord organoid; CO, cortical organoid. (E) ssGSEA heatmap showing enrichment scores for brain-region signatures. Higher scores (yellow) indicate stronger enrichment of the corresponding regional identity. (F) Immunofluorescence staining for neuronal subtype-specific markers in day 30 organoids. Scale bars, 100 μm.

To further validate regional identity, we performed bulk RNA sequencing (RNA-seq) at a later differentiation stage (day 30). Expression profiling of region-specific marker genes revealed a close resemblance to the *in vivo* medulla oblongata, while clearly separating mHBOs from previously reported brainstem organoids^24^, medullary spinal trigeminal nucleus–like organoids^25^, spinal cord organoids^26^, and cortical organoids^27^ (Figure 1D). Consistently, single-sample gene set enrichment analysis (ssGSEA) using human brain-region signature gene sets showed strong enrichment of medullary signatures in mHBOs (Figure 1E). These analyses revealed a transcriptomic landscape highly similar to that of the *in vivo* medulla oblongata and clearly distinct from previously reported organoid models.

Immunostaining at this stage of mHBOs revealed enriched populations of CHAT-positive cholinergic neurons and GABAergic interneurons compared with non-patterned or cortical organoids (Figure 1F).

### Single-cell transcriptomics reveals medullary hindbrain lineage specification

To further characterize the regional identity and lineage specification of the mHBOs, we performed single-cell RNA-seq (scRNA-seq) on 8,673 cells collected at day 50. Uniform manifold approximation and projection (UMAP) dimensionality reduction identified 22 clusters, which were further categorized into six major cell types, including radial glia, intermediate progenitor cells (IPC), excitatory neurons (ExN), inhibitory neurons (InN), astrocytes (Astro), and oligodendrocytes (Oligo) according to marker expression, as previously described^28^ (Figures 2A and 2B). Expression of HOX gene families across neuronal and progenitor clusters again confirmed a posterior (medullary) hindbrain identity (Figure 2C). To assess the regional identity of the mHBOs, we integrated the mHBO dataset with a published human fetal brain atlas.^29^ The mHBOs most closely resemble the human medulla (Figures 2D). We next examined dorsoventral (DV) patterning within mHBOs by analyzing transcription factors associated with cardinal progenitor domains. Despite the absence of exogenous Sonic hedgehog (SHH) signaling during differentiation, we observed spatially restricted expression of DV patterning markers, including EVX1 (V0), GATA3 (V2/MN), and NKX2-2 (V3/MN), indicating the specification of ventral neuronal populations (Figure 2E). By mapping these combinatorial expression profiles onto known developmental trajectories,^30–32^ we identified neuronal populations corresponding to specific medullary nuclei, including the nucleus of the solitary tract (NTS; derived from dA3 and dB1 progenitors) and the inferior olive (IO; derived from dA4 progenitors). Together, these results indicate that mHBOs recapitulate medullary neurons lineage specification by autonomously generating a broad spectrum of DV neuronal identities found in the *in vivo* medulla oblongata.

**Figure 2.**
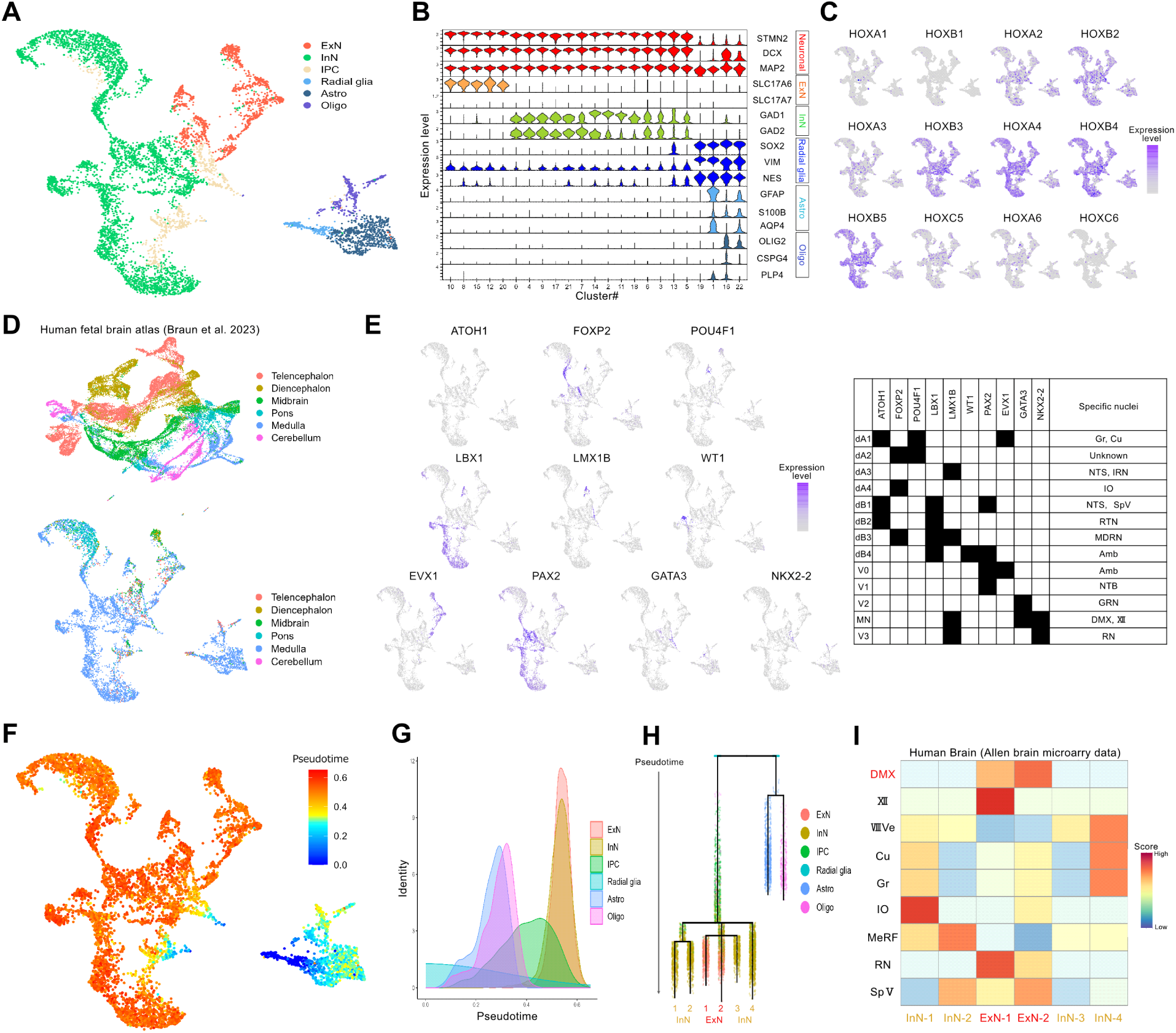
Single-cell transcriptomic analysis of mHBO. (A) UMAP embedding of single-cell RNA-seq data from day 50 mHBOs. ExN, excitatory neuron; InN, inhibitory neuron; IPC, intermediate progenitor cell; Astro, astrocyte; Oligo, oligodendrocyte. (B) Violin plots showing the expression of representative marker genes across identified clusters. (C) Feature plots showing HOX gene expression patterns in mHBOs. (D) Integration with a published human fetal brain dataset (top). UMAP embedding of mHBO cells with predicted brain-region annotations transferred from the reference atlas (bottom). (E) Feature plots showing the expression of dorsoventral patterning genes in mHBOs (left), alongside their reported expression domains in previous studies (right). Gr, gracile nucleus; Cu, cuneate nucleus; NTS, nucleus of the solitary tract; IRN, intermediate reticular zone; IO, inferior olivary complex; SpV, spinal trigeminal nucleus; RTN, retrotrapezoid nucleus; MDRN, medullary dorsal reticular nucleus; Amb, ambiguous nucleus; NTB, lateral nucleus of the trapezoid body; GRN, gigantocellular reticular nucleus; XII, hypoglossal nucleus; RN, raphe nuclei. (F) Pseudotime analysis of mHBO cells reconstructed using URD. (G) Ridgeline plots showing the pseudotime distribution of each cell type in mHBOs. (H) URD lineage tree showing branching of the excitatory neuron (ExN) lineage into two distinct trajectories. See also Figure S2. (I) AUCell heatmap showing enrichment scores for brain-region signatures derived from the Allen Brain Atlas. VIIIVe, vestibular nuclei; MeRF, medullary reticular formation.

To reconstruct developmental trajectories, we performed pseudotime analysis across all neuronal and progenitor populations. This revealed a continuous lineage progression from radial glia and IPCs toward differentiated inhibitory and excitatory neurons (Figures 2F, 2G, and S2A–C). Notably, the excitatory neuron lineage bifurcated into two distinct branches (ExN-1 and ExN-2) (Figure 2H). Differential gene expression analysis between ExN-1 and ExN-2 revealed that ExN-2 was characterized by elevated expression of TH, a previously reported marker of DMX neurons,^33^ as well as visceral motor–associated transcription factors PHOX2A and LMX1B^34^ (Figure S2D). Comparisons with previously reported DMX markers^33,35^ also revealed that ExN-2 is characterized by expression of PHOX2B, ADCYAP1, and SNCA, whereas ExN-1 showed higher levels of CHAT and SNCG (Figure S2E). To assign regional identity at the level of discrete nuclei, we compared transcriptomic signatures of neuronal subpopulations with reference data from the Allen Human Brain Atlas. Notably, ExN-2 exhibited a molecular profile that most closely resembled the human DMX, whereas ExN-1 aligned with other ventral medullary populations, including the raphe magnus and hypoglossal nuclei (Figure 2I). Based on these results, we defined ExN-2 as DMX-like neurons, confirming the successful generation of the vagal origin cells required for the brain–gut axis model.

### Generation of brain-intestinal assembloids

To reconstruct the brain–gut axis via the vagal circuit, it is essential to incorporate enteric neurons, which reside at the anatomical and functional interface between the central and peripheral nervous systems and play critical roles in gastrointestinal homeostasis and disease.^36^ However, conventional human iPSC–derived intestinal organoids (IOs) contain only sparse enteric neurons,^37,38^ limiting their utility for modeling gut–brain communication. To overcome this limitation, we adopted a dual-lineage differentiation strategy, in which enteric neural crest cells (ENCCs) and IOs were generated independently and subsequently assembled to produce enteric neuron–enriched intestinal organoids. We first differentiated hiPSCs into vagal ENCCs using a previously established protocol^39^ (Figure 3A). By day 11, spheroids contained a high proportion of SOX10-positive neural crest cells, which retained the capacity to differentiate into PRPH- and PHOX2B-positive enteric neurons, confirming successful specification of an enteric neuronal lineage (Figure 3B). In parallel, hiPSCs were differentiated into IOs following a reported protocol with minor modifications^40^ (Figure 3C). At day 4, cultures exhibited robust expression of SOX17 and FOXA2, indicating efficient induction of definitive endoderm (Figure 3D). By day 11, organoids expressed CDX2, consistent with a midgut identity from which the intestine and large portions of the colon arise, while a smaller fraction of SOX2-positive cells indicated limited foregut specification. To generate enteric neuron–enriched IOs, day 9 hindgut spheroids were combined with day 11 ENCCs. Prior to assembly, ENCCs were labeled using a lentiviral vector expressing EF1α-driven mCherry, enabling lineage tracing after integration (Figure 3E). As expected, IOs generated in isolation contained few PRPH-positive enteric neurons. In contrast, co-assembly with ENCCs resulted in robust enrichment of mCherry-positive, PRPH-positive enteric neurons, confirming efficient incorporation and differentiation of enteric neurons within the intestinal tissue (Figure 3F).

**Figure 3.**
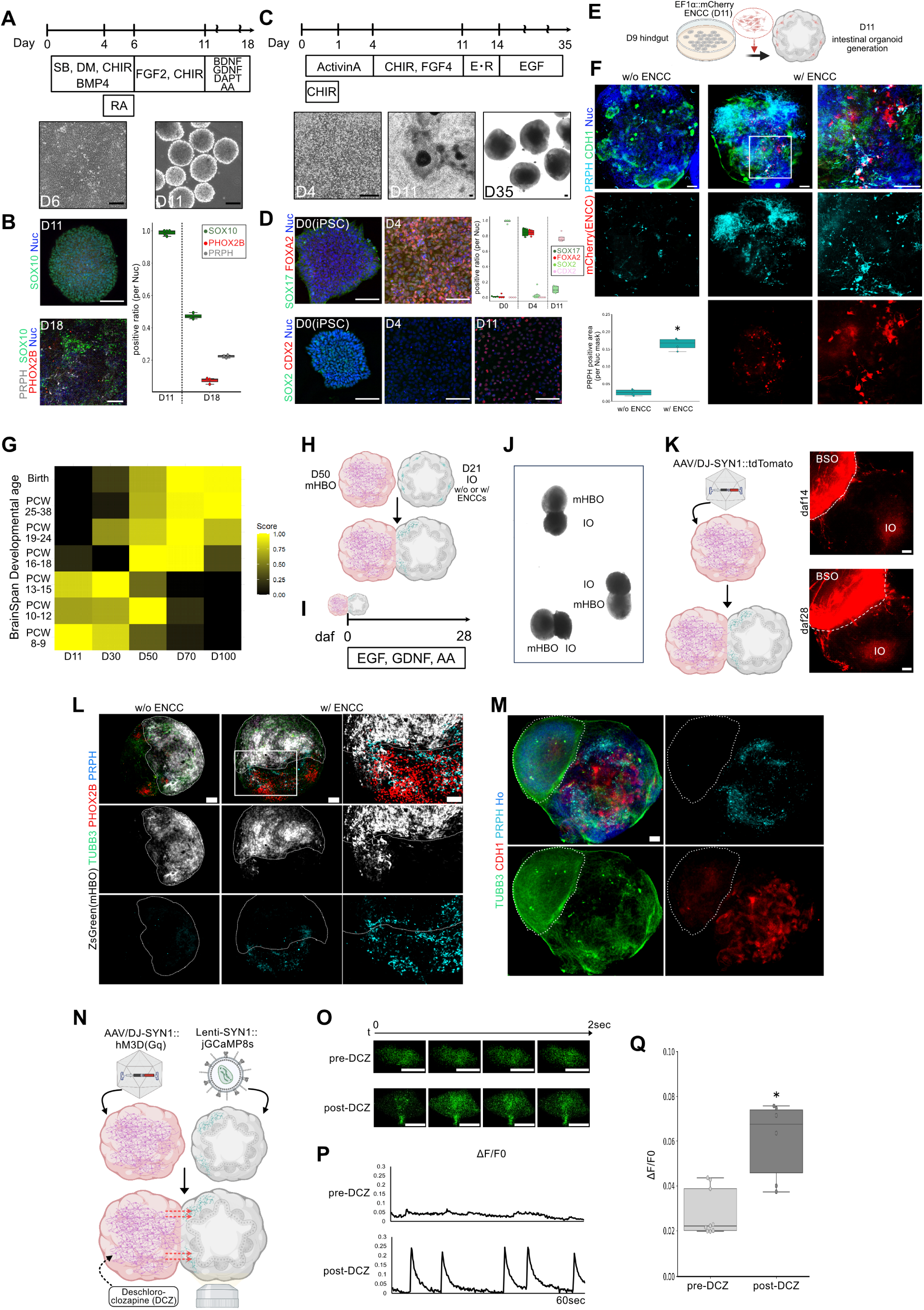
Generation of brain–intestinal assembloids. (A) Schematic of the differentiation protocol for enteric neural crest cells (ENCCs) (top). Representative brightfield images show ENCC morphology at indicated differentiation time points (bottom). Scale bar, 100 μm. (B) Immunostaining of ENCC spheroids at day 11 and differentiated neurons at day 18 (left). Quantification of marker-positive cells at each time point (right). Data are shown as box-and-whisker plots (n = 3-5). (C) Schematic of the differentiation protocol for intestinal organoids (IOs) (top). Representative brightfield images show IO morphology at indicated differentiation time points (bottom). Scale bar, 100 μm. E, EGF (epidermal growth factor); R, R-spondin 1. (D) Immunofluorescence staining of definitive endoderm (DE; day 4) and hindgut progenitors (day 11) (left). Quantification of marker-positive cells at each time point (right). Data are shown as box-and-whisker plots (n = 4-5). (E) Schematic of ENCC assembly with day 9 hindgut progenitors. For lineage tracing after assembly, ENCCs were labeled using a lentiviral vector expressing EF1α-driven mCherry. (F) Immunofluorescence staining of IOs with or without ENCC assembly at day 35. Quantification of PRPH-positive enteric neurons (bottom left). Data are shown as box-and-whisker plots (n = 4). The two-tailed Mann–Whitney test was used for comparison. ∗*P* < 0.05. (G) ssGSEA heatmap showing enrichment scores for developmental stage signatures derived from the BrainSpan dataset across sequential mHBO bulk RNA-seq samples (day 11–100). (H) Schematic of brain–intestinal assembloid (BIA) generation. (I) Schematic of the culture protocol for BIAs. Daf, days after fusion. (J) Representative bright field image of daf 14 BIA. (K) Schematic of AAV transduction into mHBOs (left). Representative fluorescence images of BIAs at daf 14 and daf 28. Scale bars, 100 μm. (L) Immunofluorescence staining of daf 14 BIAs generated with or without ENCC assembly. The dotted outline indicates the mHBO region. Scale bars, 100 μm. (M) Immunofluorescence staining of a daf 28 BIA. The dotted outline indicates the mHBO region Scale bars, 100 µm. (N) Schematic of the calcium imaging experimental setup. (O) Representative images showing spontaneous (pre-DCZ) and DCZ-induced (post-DCZ) calcium transients over a 2-second interval. Scale bars, 10 μm. (P) Representative ΔF/F₀ traces before and after DCZ treatment. (Q) Quantification of ΔF/F₀. Data are shown as box-and-whisker plots (n = 6–9). Unpaired two-tailed Student’s t-test was used for comparison. \**P* < 0.05.

To generate brain–intestinal assembloids (BIAs), we first determined the optimal developmental timing for assembly. In humans, vagal nerve projections toward peripheral targets are initiated around post-conception week (PCW) 8,^41,42^ followed by progressive target innervation and functional maturation. Consistently, ssGSEA-based comparison of sequential hindbrain organoids with the Allen Human Brain Developmental Atlas indicated that day 50 organoids transcriptionally correspond to PCW 10–12, a developmental window associated with ongoing vagal target innervation and emerging functional connectivity, whereas later culture stages align with more advanced developmental periods (PCW 19–24) (Figure 3G). Based on this analysis, day 50 was selected to model a developmental stage permissive for establishing functional brain–gut connectivity. Accordingly, day 50 mHBOs were fused with day 21 IOs either with or without ENCCs (Figures 3H–J). To confirm sequential neuronal innervation within the BIAs, we employed adeno-associated virus (AAV) vectors expressing Synapsin-1 promoter–driven tdTomato (AAV-hSYN1::tdTomato) (Figure 3K). Fluorescent tracing revealed progressive axonal extension from the medullary hindbrain organoid toward the intestinal compartment following assembly. Immunostaining further confirmed that pre-assembly incorporation of ENCCs enabled robust neuronal innervation from the mHBOs and promoted the emergence of PRPH-positive enteric neurons localized at the interface between brain and gut tissues (Figure 3L). In addition, we also confirmed coordinated organization of the three major components, namely brainstem neurons (TUBB3 positive), enteric neurons (PRPH positive), and intestinal epithelial cells (CDH1 positive), within a single integrated BlAs (Figure 3M).

To validate functional connectivity between mHBOs and enteric neurons, we next examined activity-dependent neuronal activation. We selectively activated hindbrain neurons using a chemogenetic approach and assessed downstream responses in enteric neurons. mHBOs were infected with AAV-hSYN1::hM3D(Gq), and assembled with IOs and matured until day 21 after fusion (daf 21). Activation of hM3D(Gq) was achieved using deschloroclozapine (DCZ), a highly potent and selective agonist.^43^ Immunostaining for c-Fos, an immediate early gene rapidly induced by neuronal depolarization and synaptic activity,^44^ revealed that neither mock-treated BIAs nor hM3D(Gq)-expressing BIAs in the absence of DCZ showed appreciable c-Fos induction. In contrast, DCZ treatment of hM3D(Gq)-expressing BIAs resulted in a significant increase in c-Fos–positive enteric neurons within the intestinal compartment (Figures S3A and S3B), indicating activity-dependent signal transmission from brainstem neurons to enteric neurons. We next performed live calcium imaging to directly visualize functional connectivity between the two compartments (Figure 3N). Baseline spontaneous calcium oscillations in enteric neurons were rarely observed prior to stimulation. Upon DCZ application, selective activation of hindbrain neurons triggered a robust increase in calcium transients (ΔF/F₀) within enteric neurons of the IOs (Figures 3O and 3P). This increase in firing frequency and amplitude provided direct evidence of functional signal propagation from brainstem neurons to enteric neurons across the BIAs (Figure 3Q). Together, these findings indicate that the BIAs establish not only anatomically organized vagal-like innervation but also functional connectivity between medullary hindbrain neurons and enteric neurons, recapitulating a key feature of the human brain–gut axis *in vitro*.

### Identification of aberrant cell–cell signaling underlying brain-to-gut α-synuclein propagation

To model pathological α-syn transmission along the brain–gut axis, we established a system to selectively introduce α-syn into the brain compartment of the BIAs. To distinguish exogenously introduced α-syn from endogenous protein and to trace donor and recipient cells, we generated an AAV vector encoding human SNCA (α-synuclein) fused to a FLAG epitope, followed by a 2A peptide and tdTomato reporter (AAV-hSNCA/FLAG-2A-tdTomato) (Figure 4A). Efficient α-syn expression was confirmed by robust tdTomato fluorescence and FLAG immunostaining in transduced cells (Figure 4B). To specifically model brain-to-gut α-syn propagation, the AAV was introduced into mHBOs prior to assembly, after which the labeled mHBOs were fused with IOs containing enteric neurons (Figure 4C). Daf 21 assembloids showed that exogenous α-syn was deposited in the enteric neural filaments, indicating successful recapitulation of brain-to-gut α-syn propagation (Figure 4D).

**Figure 4.**
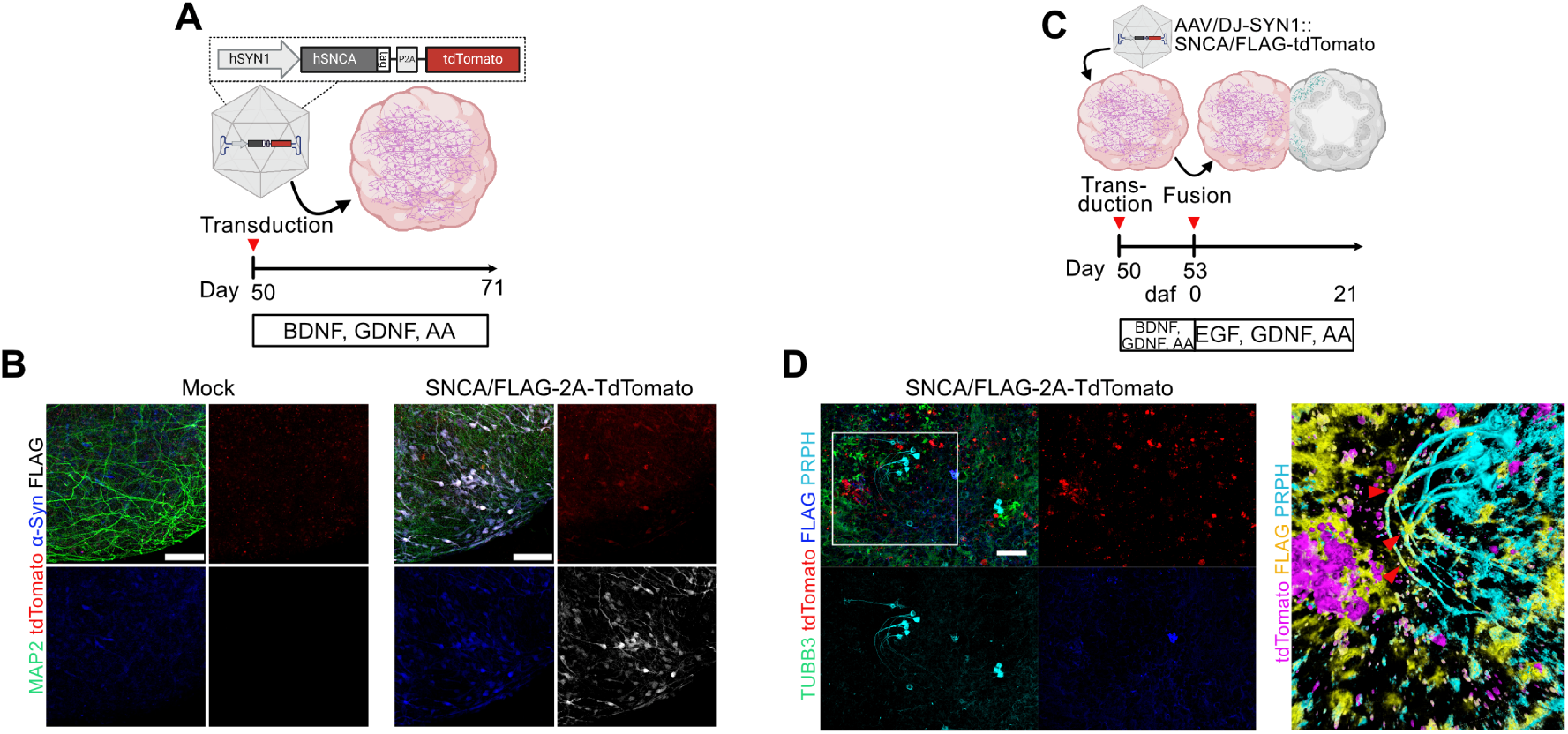
Recapitulation of brain-to-gut α-synuclein propagation in BIAs. (A) Schematic of AAV-mediated expression of FLAG-tagged human α-synuclein (α-syn). (B) Immunofluorescence staining of day 71 mHBOs transduced with AAV-SNCA. Scale bars, 100 μm. (C) Schematic of the brain-to-gut α-syn propagation assay using BIAs. (D) Immunofluorescence staining of daf 21 BIAs following AAV-SNCA transduction. A three-dimensional rendering of the boxed region is shown at right. Arrows indicate FLAG-positive α-syn deposition along enteric neuronal fibers. Scale bars, 100 μm.

To dissect the molecular mechanisms underlying brain-to-gut α-syn propagation, we performed ligand–receptor interaction analysis using integrated single-cell transcriptomic data. We generated a combined single-cell RNA-seq dataset comprising mHBOs, BIAs under mock conditions, and BIAs with AAV-mediated SNCA overexpression (Figure 5A). Cell types were annotated based on established marker genes, following previous studies^38^ (Figure 5B and 5C). The overall cellular composition was highly comparable between mock and SNCA-overexpressing assembloids, indicating that α-syn overexpression did not perturb cell-type specification (Figure 5D). As expected, SNCA mRNA levels were selectively increased in the hindbrain compartment of SNCA-overexpressing assembloids, confirming compartment-restricted α-syn expression (Figure 5E). We next performed ligand–receptor interaction analysis using CellChat v2.^45^ Under mock conditions, the BIAs exhibited extensive intercellular communication among neuronal and non-neuronal populations (Figure S4A). Notably, physiologically relevant signaling pathways were preserved, including glutamatergic signaling among neuronal populations, including enteric neurons, consistent with long-range neuronal communication,^46^ as well as WNT signaling from intestinal mesenchyme to epithelial compartments, reflecting niche-supporting interactions in the gut^47^ (Figure S4B). These results indicate that the BIAs recapitulate key aspects of physiological cell–cell communication.

**Figure 5.**
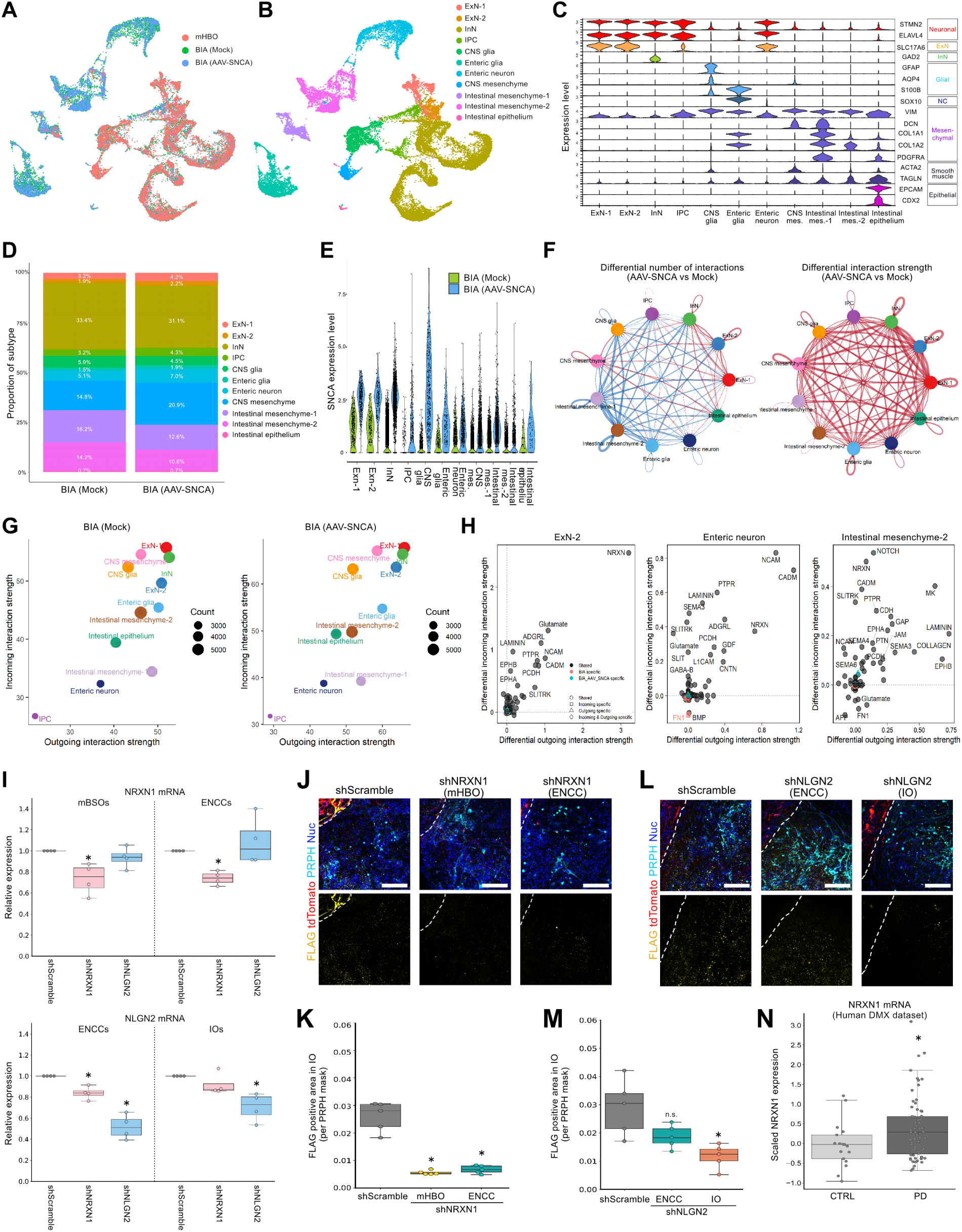
NRXN1–NLGN2 signaling mediates brain-to-gut α-syn propagation. (A) Integrated UMAP embedding of scRNA-seq data from day 71 mHBOs and daf 21 BIAs with AAV-SNCA transduction. (B) UMAP visualization of annotated cell types. (C) Violin plots showing the expression of representative marker genes across identified cell types. (D) Comparison of cell-type composition between mock and SNCA-overexpressing BIAs. (E) Violin plots showing SNCA expression across cell types. (F) Summary of global cell–cell interaction differences across all cell types between mock and SNCA–overexpressing BIAs. Red indicates increased interaction strength, whereas blue edges indicate decreased interaction strength in SNCA-overexpressing BIAs relative to mock. Thickness reflects the magnitude of change. (G) Differential interaction strength matrix at the cell-type level comparing mock and SNCA-overexpressing BIAs. (H) Differential signaling pathway interaction strengths in ExN-2, enteric neurons, and intestinal mesenchyme-2. (I) Knockdown efficiency of NRXN1 or NLGN2 assessed by RT–qPCR in mHBOs, IOs (without ENCC assembly), and ENCC-derived neurons (2D culture). Data are shown as box-and-whisker plots (n = 4). The Steel test was used for comparison. \**P* < 0.05 compared with shScramble. (J) Immunofluorescence staining of daf 21 BIAs with or without shNRXN1. The dotted line indicates the boundary between the mHBO and IO regions. Scale bars, 100 μm. (K) Quantification of FLAG-positive area within PRPH-positive enteric neuronal fibers in (J). Data are shown as box-and-whisker plots (n = 5). The Steel test was used for comparison. \**P* < 0.05 compared with shScramble. (L) Immunofluorescence staining of daf 21 BIAs with or without shNLGN2. The dotted line indicates the boundary between the mHBO and IO regions. Scale bars, 100 μm. (M) Quantification of FLAG-positive area within PRPH-positive enteric neuronal fibers in (L). Data are shown as box-and-whisker plots (n = 5). The Steel test was used for comparison. \**P* < 0.05 compared with shScramble. n.s., not significant. (N) NRXN1 expression in excitatory neurons of DMX in control and PD samples. Data are shown as box-and-whisker plots (n = 17 controls, n = 67 PD). Unpaired two-tailed t-test was used for comparison. \**P* < 0.05 compared with control. See also Figure S8.

To identify aberrant signaling associated with α-syn propagation, we compared ligand–receptor interaction strength between mock and SNCA-overexpressing assembloids. Although the total number of interactions varied, global interaction strength was increased across nearly all cell-type pairs in the SNCA-overexpression condition (Figure 5F), suggesting widespread remodeling of intercellular communication. Among all populations, the most pronounced changes were observed between ExN-2 (DMX-like excitatory neurons) and intestinal mesenchyme-2 (Figure 5G). Given that DMX-like neurons represent the presumptive source of α-syn and enteric neurons represent the recipient population, we examined outgoing signals from ExN-2 neurons. Family-level analysis identified neurexin (NRXN) signaling as the most strongly upregulated outgoing pathway from ExN-2 neurons in SNCA-overexpressing assembloids (Figure 5H). Interestingly, NRXN signaling was not among the top altered incoming signals in enteric neurons, but instead emerged as a dominant interaction in intestinal mesenchyme-2, suggesting a non-canonical route of signal relay. At the ligand–receptor pair level, NRXN1–NLGN2 exhibited the highest communication probability among NRXN1-associated ligand-receptor pairs, particularly along the ExN-2 and intestinal mesenchyme-2 under SNCA-overexpression conditions (Figure S4C). Neurexins are well-established presynaptic adhesion molecules critical for synapse organization.^48^ However, expression of other synaptic markers was not globally altered in SNCA-overexpressing assembloids (Figure S4D), arguing against a simple increase in synaptogenesis as the mechanism of propagation. Notably, α-syn fibrils have been reported to directly bind presynaptic NRXN1,^49^ whereas NLGN2 has not been previously implicated in α-syn binding, and its physiological role in intestinal mesenchyme remains poorly understood. To validate the expression pattern of NLGN2 in the human gut, we reanalyzed a publicly available human gut cell atlas^50^ (Figure S5A and S5B). This analysis confirmed that NLGN2 is specifically expressed in mesenchymal populations, whereas NRXN1 is restricted to neuronal clusters, supporting the spatial plausibility of this ligand–receptor interaction *in vivo* (Figure S5C and S5D).

To functionally validate the role of NRXN1–NLGN2 signaling in α-syn propagation, we performed knockdown experiments using lentiviral shRNA vectors introduced prior to assembloid assembly. Knockdown efficiency was confirmed by RT-qPCR (Figure 5I). Knockdown of NRXN1 in either mHBOs or enteric neurons significantly reduced FLAG-positive α-syn deposition in enteric neuronal fibers, indicating effective attenuation of α-syn propagation (Figures 5J and 5K). These results are consistent with previous reports identifying NRXN1 as a neuronal receptor for α-syn.^49^ In contrast, NLGN2 knockdown in enteric neurons did not significantly affect α-syn deposition. Although shNRXN1 knockdown led to changes in NLGN2 expression within ENCCs (Figure 5I), direct perturbation of NLGN2 in enteric neurons did not affect α-syn propagation, indicating that NLGN2 activity in enteric neurons is not a major determinant of α-syn propagation. However, NLGN2 knockdown in IOs markedly reduced α-syn accumulation in enteric neurons (Figures 5L and 5M), indicating that intestinal mesenchymal NLGN2 acts as a critical mediator of brain-to-gut α-syn propagation. NRXN1 knockdown did not sufficiently attenuate brain–to–brain α–syn propagation (Figures S6A–C). We further tested whether the NRXN signal is also involved in α-syn aggregate propagation, which is considered more pathogenic than synuclein monomer.^51^ We employed a recently reported self-assembling α-syn system, which induces intracellular α-syn aggregation via doxycycline-inducible expression of α-synuclein fused to a self-assembling protein.^52^ This system successfully generated aggregated α-syn in both HEK293T cells and mHBO (Figures S6D-F). This system also achieved brain–gut synuclein propagation, and NRXN1 knockdown in mHBOs significantly attenuated the propagation of aggregated V5–tagged α–syn. These results indicate that NRXN1 is also involved in the pathological aggregated α-syn transmission (Figures S6G-I). Finally, to assess the clinical relevance of these findings, we reanalyzed a publicly available multi-region single-nucleus RNA-seq atlas of PD brains.^53^ Focusing on the DMX across four brain banks, we isolated excitatory neuronal populations based on marker gene expression (Figure S7A-E). After quality-filtering, the dataset comprised 4,908 cells from 17 healthy individuals and 67 PD patients, with comparable age and sex distributions (Figure S7A). Notably, NRXN1 expression was significantly elevated in PD patients compared with healthy controls, whereas NRXN2 and NRXN3 showed no significant differences (Figure 5N, S7F and S7G). Together, these results identify NRXN1–NLGN2-mediated aberrant cell–cell signaling as a previously unrecognized mechanism facilitating brain-to-gut α-synuclein propagation.

## DISCUSSION

In this study, we established a human iPSC-derived BIA that reconstructs a physiologically relevant vagal circuit connecting medullary hindbrain neurons and enteric neurons, and we subsequently leveraged this platform to dissect the cellular and molecular mechanisms underlying brain-to-gut α-syn propagation in PD. By integrating regionally patterned brainstem organoids, assembloid technology, single-cell transcriptomics, and genetic perturbation, our work provides a human-specific framework for understanding how pathological signals traverse the brain–gut axis.

Vagal motor neurons represent a critical gateway of the gut–brain axis, serving as the principal efferent output from the brain to visceral organs. Neurons in the DMX regulate a wide range of autonomic functions, including gastrointestinal motility and secretion, heart rate modulation, swallowing, and aspects of vocalization and airway control, through parasympathetic innervation of thoracic and abdominal organs.^54^ In addition to these physiological roles, accumulating pathological and epidemiological evidence positions the DMX as one of the earliest affected brain regions in Parkinson’s disease, underscoring its importance as a disease-relevant neuronal population.^6,7,9^ Despite this central role, faithful *in vitro* modeling of human vagal motor neurons remains limited, as existing studies have primarily generated peripheral vagal parasympathetic neurons,^55,56^ and no reports to date have successfully derived presynaptic DMX-like preganglionic neurons from human stem cells. Over the past decade, both unguided cerebral organoids and region-specific brain organoids have been developed.^57–59^ However, although several hindbrain organoid protocols have been reported,^24,25,60^ the generation of medullary hindbrain organoids that reproducibly recapitulate vagal motor neurons with nucleus-level resolution has remained largely unexplored. To address this challenge, we optimized a directed differentiation protocol to generate mHBOs and validated their regional identity by comparison with transcriptomic references from the human developing brain atlas.^29^ While this approach achieved robust medullary patterning, only a subset of neurons acquired a DMX-like identity, highlighting a fundamental and previously underappreciated challenge. Notably, our mHBOs autonomously generated a broad spectrum of dorsoventral identities, providing a suitable framework to interrogate neuronal fate decisions rather than imposing overly restrictive patterning cues. Looking forward, generating nucleus-specific DMX organoids or multi-nucleus medullary organoids incorporating additional vagal-related nuclei such as the nucleus tractus solitarius will be essential to model higher-order autonomic circuits, including vagovagal reflexes and bidirectional gut–brain communication. Achieving this goal will likely require precise control of spatiotemporal gene regulation through strategies such as refined morphogen gradient engineering, CRISPR-based transcriptional modulation, and the assembly of highly patterned, nucleus-level organoids.

Although mounting evidence implicates the brain–gut axis in PD pathogenesis, experimental investigation of this circuit has been severely constrained by the lack of appropriate human *in vitro* models. Recent efforts have begun to address this gap using microfluidic platforms to connect human gut and brain cells *in vitro*.^16^ While these systems elegantly enable bidirectional signaling and controlled microenvironments, they predominantly rely on planar, two-dimensional architectures. In this study, we overcome these limitations by reconstructing a human gut–brain axis *in vitro* using a three-dimensional assembloid strategy. Assembloid technologies have rapidly expanded beyond the fusion of multiple brain regions to encompass increasingly complex inter-organ circuits.^61,62^ Our brain–gut assembloid faithfully recapitulates sequential neuronal innervation from medullary brainstem neurons to enteric neurons; the spatial organization of the medulla oblongata, enteric nervous system, and intestinal epithelial compartments; and functional signal transmission from brainstem neurons to enteric neurons. These properties distinguish our system from prior organ-on-chip or planar co-culture approaches and establish a unique experimental platform to investigate human-specific mechanisms of gut–brain communication.

Using the BIAs, we directly demonstrated descending propagation of α-syn from medullary neurons to enteric neuronal filaments. Integrated single-cell transcriptomics and ligand–receptor interaction analysis uncovered widespread remodeling of intercellular communication in response to α-syn overexpression, with particularly pronounced changes involving DMX-like excitatory neurons and intestinal mesenchymal cells. Among these, NRXN1–NLGN2 signaling emerged as a dominant and previously unrecognized pathway associated with α-syn propagation. Neurexins are classically viewed as presynaptic adhesion molecules involved in synapse organization.^48^ However, our data suggest that their contribution in this context extends beyond canonical synaptogenesis. Although direct visualization of synaptic activity changes induced by α-syn overexpression or genetic perturbation remains unverified, global expression of synaptic markers was not broadly altered, arguing against a simple enhancement of synapse formation as the underlying mechanism. Consistent with a non-canonical role, NLGN2 expression was primarily confined to intestinal mesenchymal cells *in vivo*. While comparable NLGN2 expression was detectable in enteric neurons in BIAs, knockdown of NLGN2 in enteric neurons did not affect α-syn propagation, whereas suppression of NLGN2 in the intestinal compartment significantly attenuated propagation. This pattern indicates that mesenchymal NLGN2, rather than neuronal NLGN2, is functionally relevant and suggests that NLGN2-mediated mesenchymal processing plays a critical role in brain-to-gut α-syn transmission. In contrast, NRXN1 knockdown in either medullary neurons or enteric neurons within the BIA setting reduced α-syn propagation, indicating a requirement for neuronal NRXN1 on both the sending and receiving sides of the brain–gut interface. Notably, this dependency was not observed in mHBO–mHBO assembloids, where NRXN1 knockdown in either the sender or the receiver mHBO failed to attenuate transmission, highlighting a context-dependent function of NRXN1. In this framework, intestinal mesenchyme acts as an essential intermediate that enables efficient transfer to enteric neurons, whereas in purely neuronal contexts lacking this mesenchymal relay, alternative receptors or uptake pathways likely predominate. Supporting the clinical relevance of this pathway, reanalysis of single-nucleus RNA-seq data from PD patient brains revealed selective upregulation of NRXN1 in DMX excitatory neurons, but not of other neurexin family members, suggesting that elevated NRXN1 expression may increase susceptibility to brain-to-gut α-syn propagation in PD. Further mechanistic studies will be required to clarify how NRXN1–NLGN2 signaling facilitates α-syn propagation at the molecular and cellular levels, including whether it regulates ligand binding, vesicular uptake, or trans-tissue trafficking dynamics.

## Limitations of the study

Although our mHBOs recapitulate key features of human medulla development, they do not selectively enrich for DMX neurons. Consequently, vagal motor neuron–specific neurotransmission and its direct effects on gut physiological functions could not be fully assessed. While organoid developmental timing was aligned with human post-conceptional stages using transcriptomic references, *in vitro* cultures do not capture aging-associated processes, which are central to PD pathogenesis.

In addition, although we identified the NRXN1–NLGN2 signaling axis as a mediator of brain-to-gut α-syn propagation, NRXN1 and NLGN2 are also implicated in neurodevelopmental disorders,^63,64^ and the functional consequences of postnatal modulation of this pathway in the context of PD remain unclear. Further *in vitro* and *in vivo* studies will be needed to evaluate how perturbation of this axis affects functional connectivity and circuit integrity, as well as to elucidate the molecular mechanisms by which intestinal NLGN2 facilitates α-syn propagation. Moreover, while attenuation of α-syn propagation was achieved in the assembloid system, we did not directly assess gut functional phenotypes such as altered motility or constipation, in part because of the limited physiological maturity of IOs. Finally, our model primarily captures a brain-to-gut propagation route, but extending this framework to include retrograde transmission and additional autonomic circuits will be essential for modeling the bidirectional brain–gut axis in PD.

## RESOURCE AVAILABILITY

### Lead contact

Requests for further information and resources should be directed to and will be fulfilled by the lead contact, Wado Akamatsu.

### Materials availability

The viral vector plasmids are available from VectorBuilder. The reference codes are provided in the key resources table.

### Data and code availability

Sequencing data have been deposited in ArrayExpress and will be made publicly available upon publication. The accession numbers for the datasets analyzed in this study will be listed in the Key Resources Table at the time of publication. This study does not report original algorithms.

## ACKNOWLEDGMENTS

This study was supported by the Ministry of Education, Culture, Sports, Science and Technology (MEXT)–supported Programs for the Strategic Research Foundation at Private Universities (S1411007), and by Grant-in-Aid for Scientific Research (24K23249 to A.Y. and 23K06934 to K.I.) from the Japan Society for the Promotion of Science (JSPS). We appreciate Editage (www.editage.com) for its English-language editing service.

## AUTHOR CONTRIBUTIONS

Conceptualization, A.Y. and W.A.; methodology, A.Y.; Investigation, A.Y.; writing—original draft, A.Y.; writing—review & editing, K.I. and W.A.; funding acquisition, A.Y., K.I., and W.A. All authors have reviewed and approved the manuscript.

---

**Figure S1.**
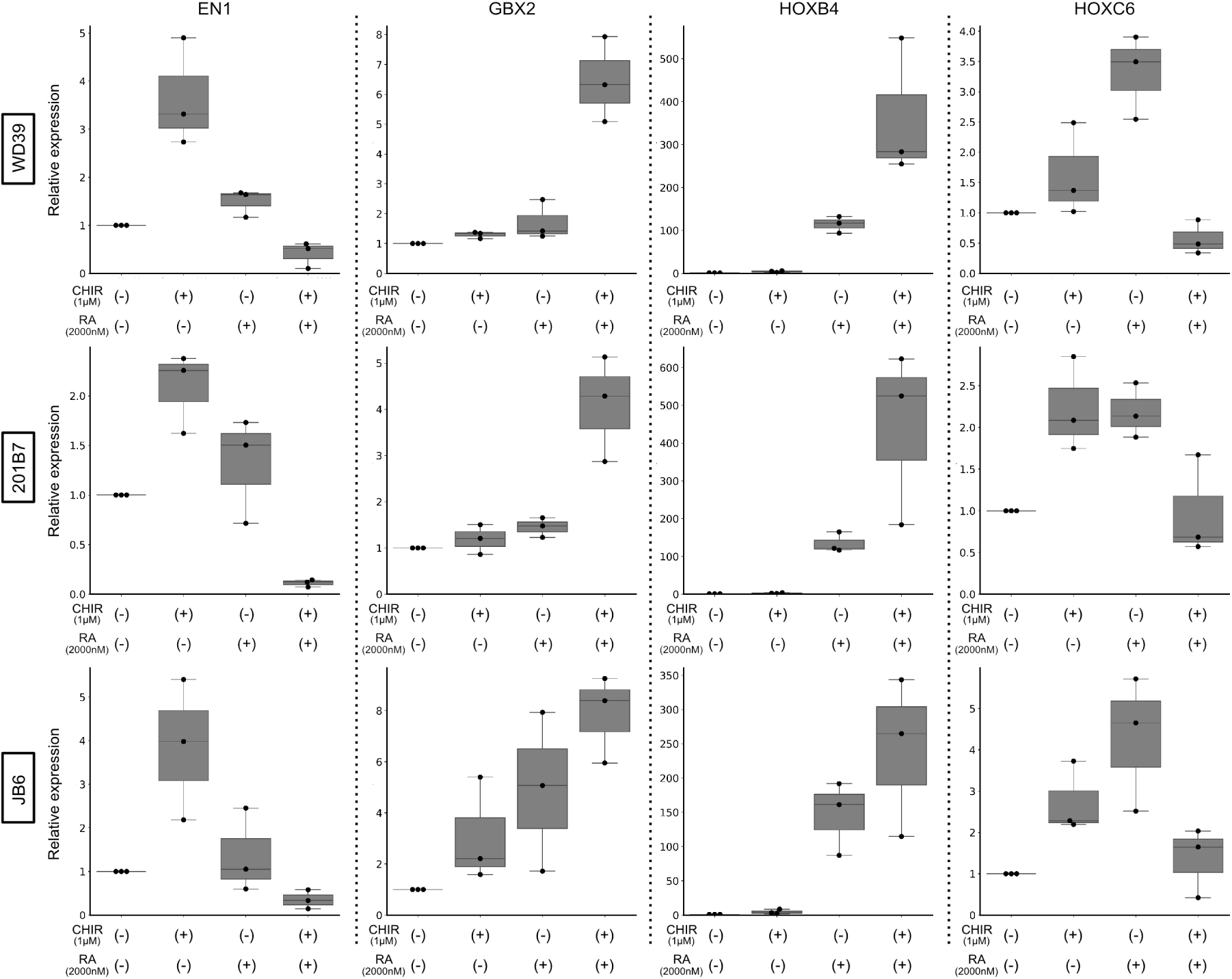
Reproducibility of the mHBO differentiation protocol across hiPSC lines, related to Figure 1. Box plots showing RT–qPCR expression of region-specific marker genes in day 11 organoids derived from three independent healthy control hiPSC lines. Data presented in the main figures were obtained using the WD39 line. Data are shown as box-and-whisker plots (n = 3 independent differentiations).

**Figure S2.**
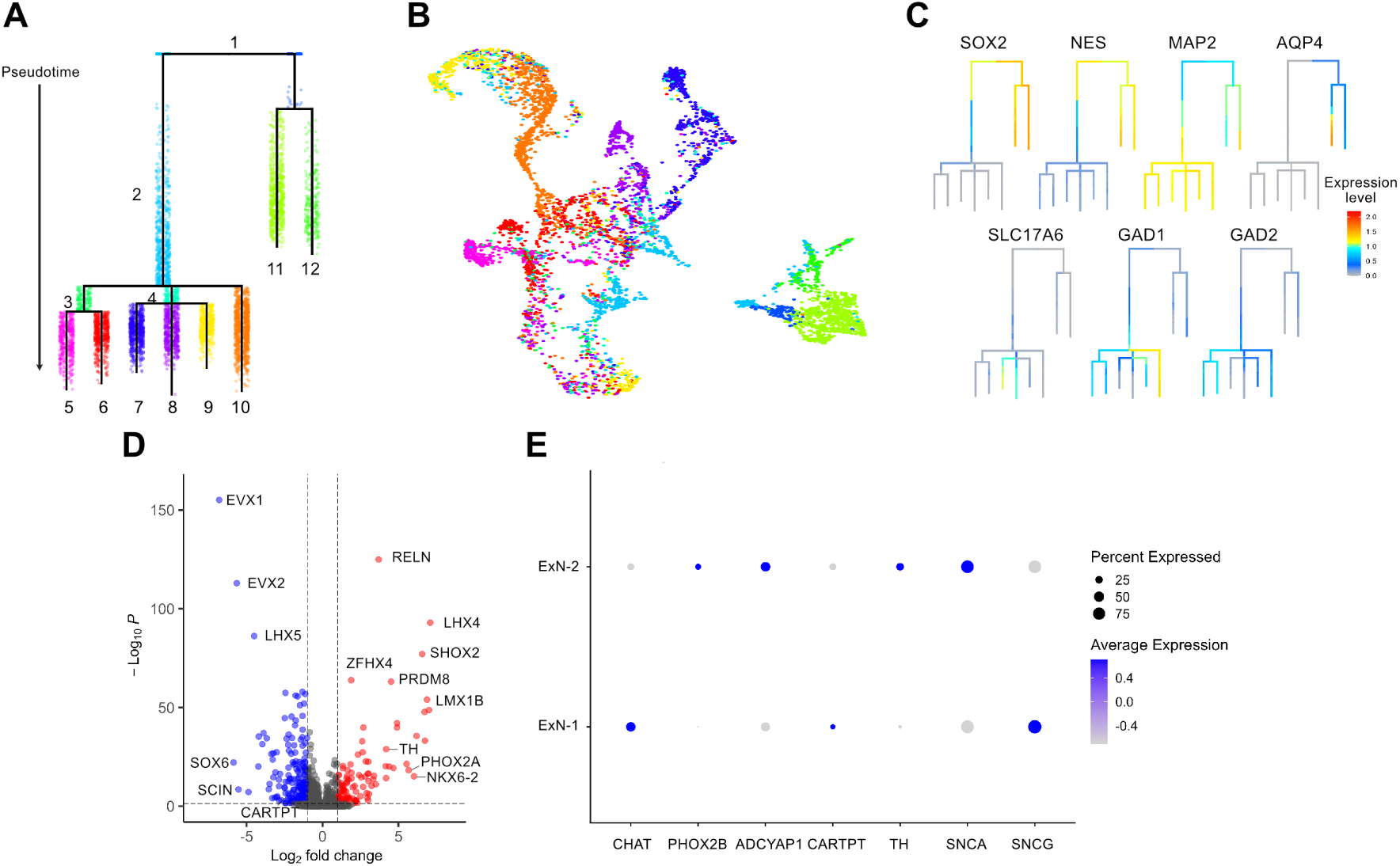
Validation of cell-type annotation and pseudotime analysis, related to Figure 2. (A) URD lineage tree illustrating developmental segments along pseudotime. (B) UMAP embedding of mHBO scRNA-seq data, colored by URD segments corresponding to those shown in (A). (C) Expression of representative marker genes mapped onto the URD lineage tree. (D) Volcano plot showing differential gene expression between ExN-2 and ExN-1 clusters. Selected genes are labeled. Red and blue dots indicate significantly up- or downregulated genes, respectively (|log₂ fold change| > 1, −log₁₀ P > 0.05). (E) Dot plot showing the expression of previously reported DMX markers. Gene expression values were scaled before visualization. Dot size represents the fraction of expressing cells, and color intensity indicates scaled expression levels.

**Figure S3.**
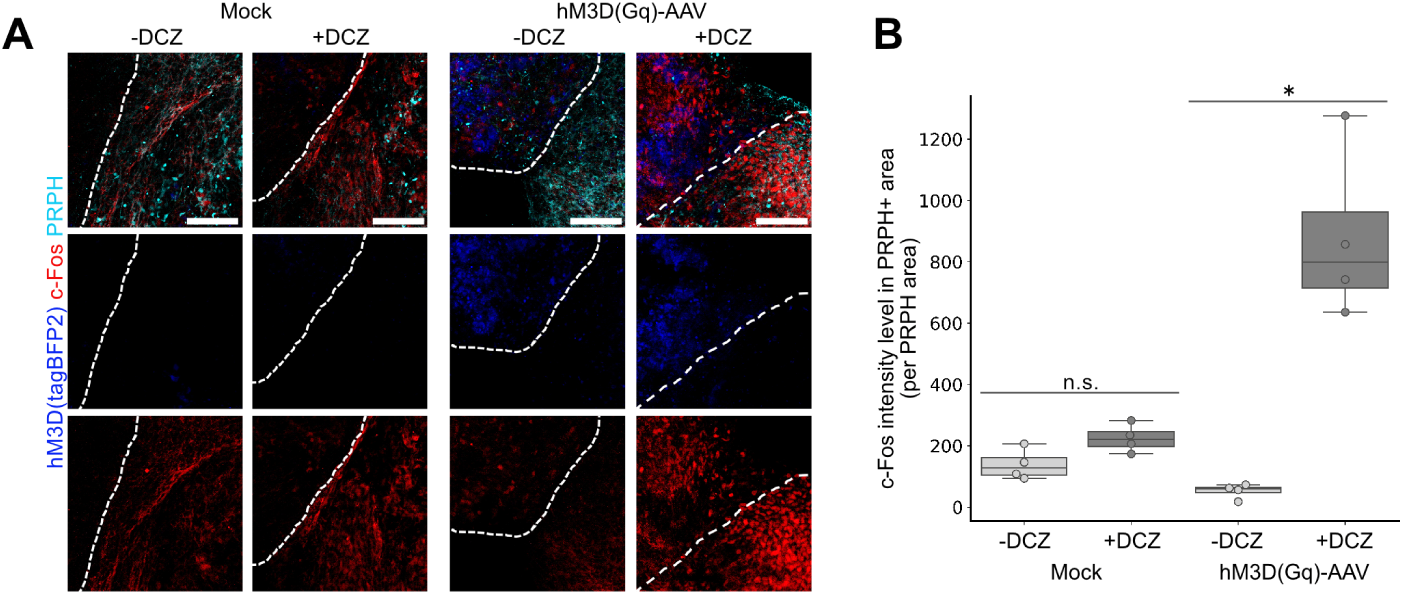
Chemogenetic activation induces c-Fos expression in brain–gut assembloids, related to Figure 3. (A) Immunofluorescence staining for c-Fos in BIAs with or without chemogenetic activation of mHBOs. The dotted line indicates the boundary between the mHBO and IO regions. Scale bar, 100 µm. (B) Quantification of c-Fos intensity within IO regions. Data are shown as box-and-whisker plots (n = 4). The two–tailed Mann–Whitney test was used for comparison. ∗*P* < 0.05. n.s., not significant.

**Figure S4.**
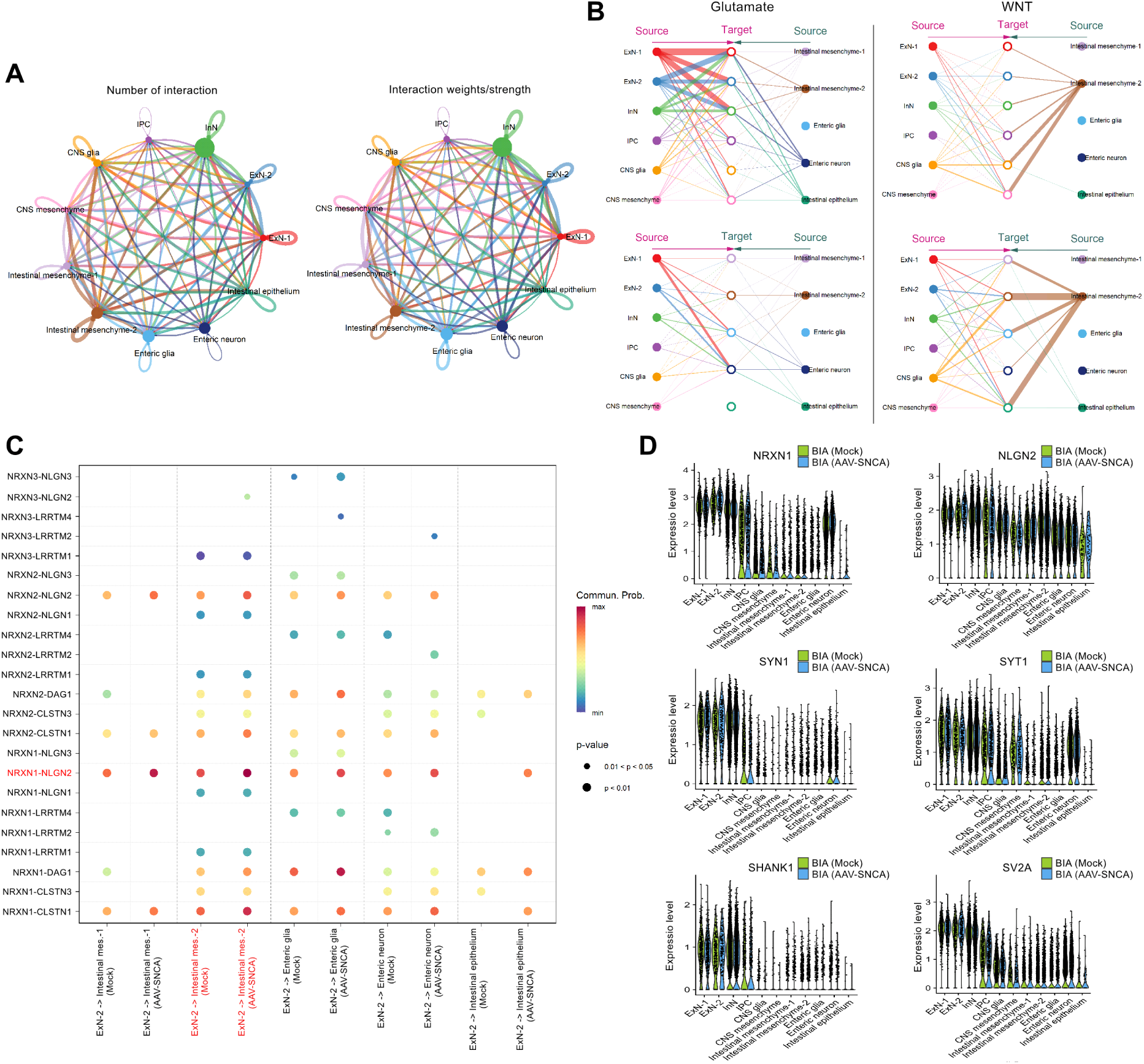
Cell–cell communication analysis and validation of NRXN signaling, related to Figure 5. (A) Summary of global cell–cell interactions among all cell types. (B) Network plots illustrating cell–cell interaction strengths for Glutamate (left) and WNT (right) signaling pathways. Line thickness represents the inferred communication strength between source and target cell populations. Target node colors correspond to the source cell types. (C) Dot plot showing predicted ligand-receptor interaction related to NRXN signaling between ExN-2 and various target cells under Mock and AAV-SNCA conditions. Dot color represents the communication probability, and dot size indicates the level of statistical significance. (D) Violin plots showing expression of NRXN1, NLGN2, and synapse-associated marker genes across cell types under Mock and AAV-SNCA conditions

**Figure S5.**
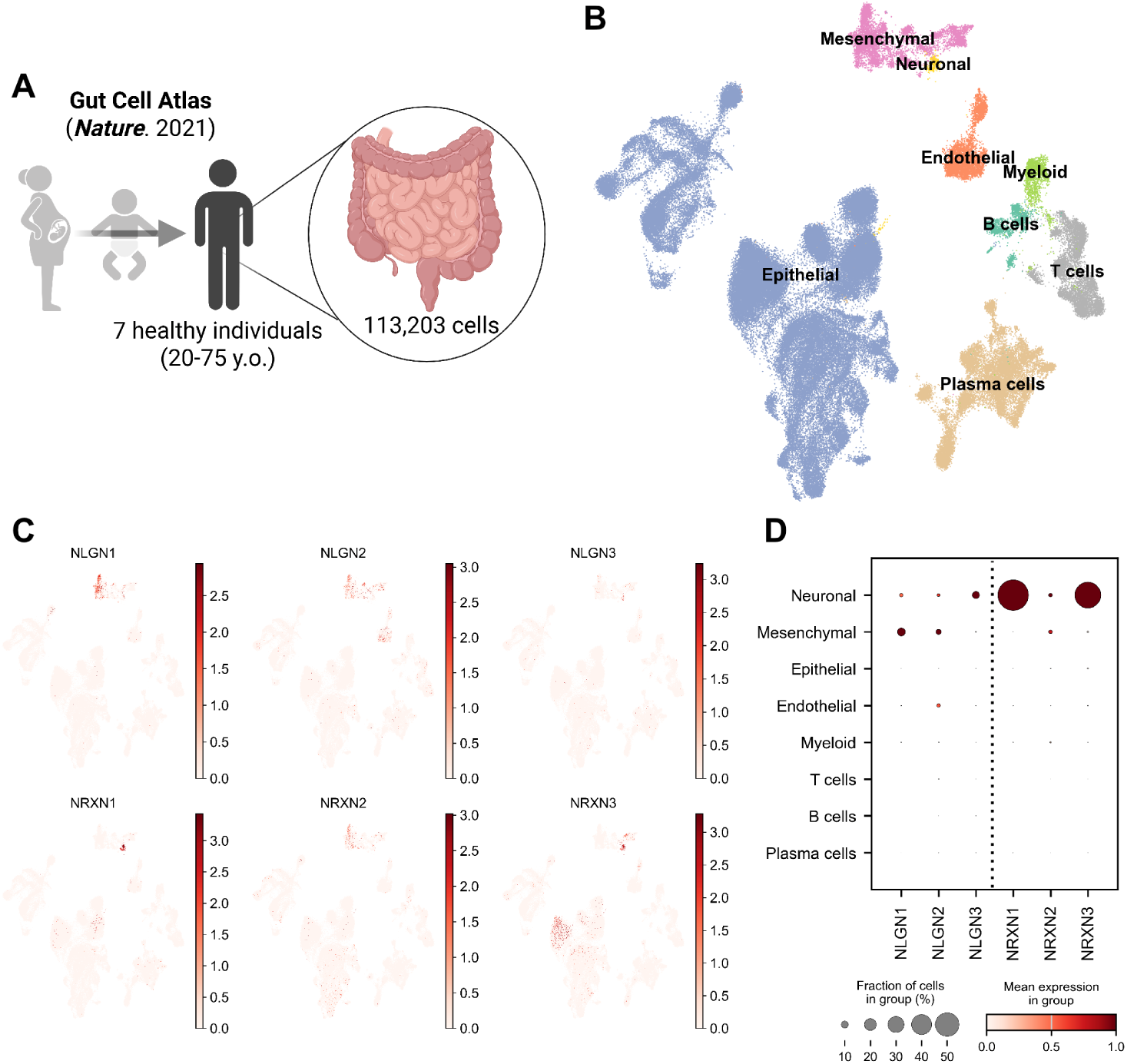
Reanalysis of the human gut cell atlas dataset, related to Figure 5. (A) Schematic of the reanalysis workflow for the human gut cell atlas dataset. Seven healthy individuals (ages 20–75 years) were selected from the original dataset for analysis. (B) UMAP embedding of the reanalyzed gut cell atlas dataset with cell-type annotations transferred from the original study. (C) Feature plots showing the expression of NRXN and NLGN family members. (D) Dot plot summarizing NRXN and NLGN expression across gut cell types. Dot size represents the fraction of expressing cells, and color intensity indicates expression levels.

**Figure S6.**
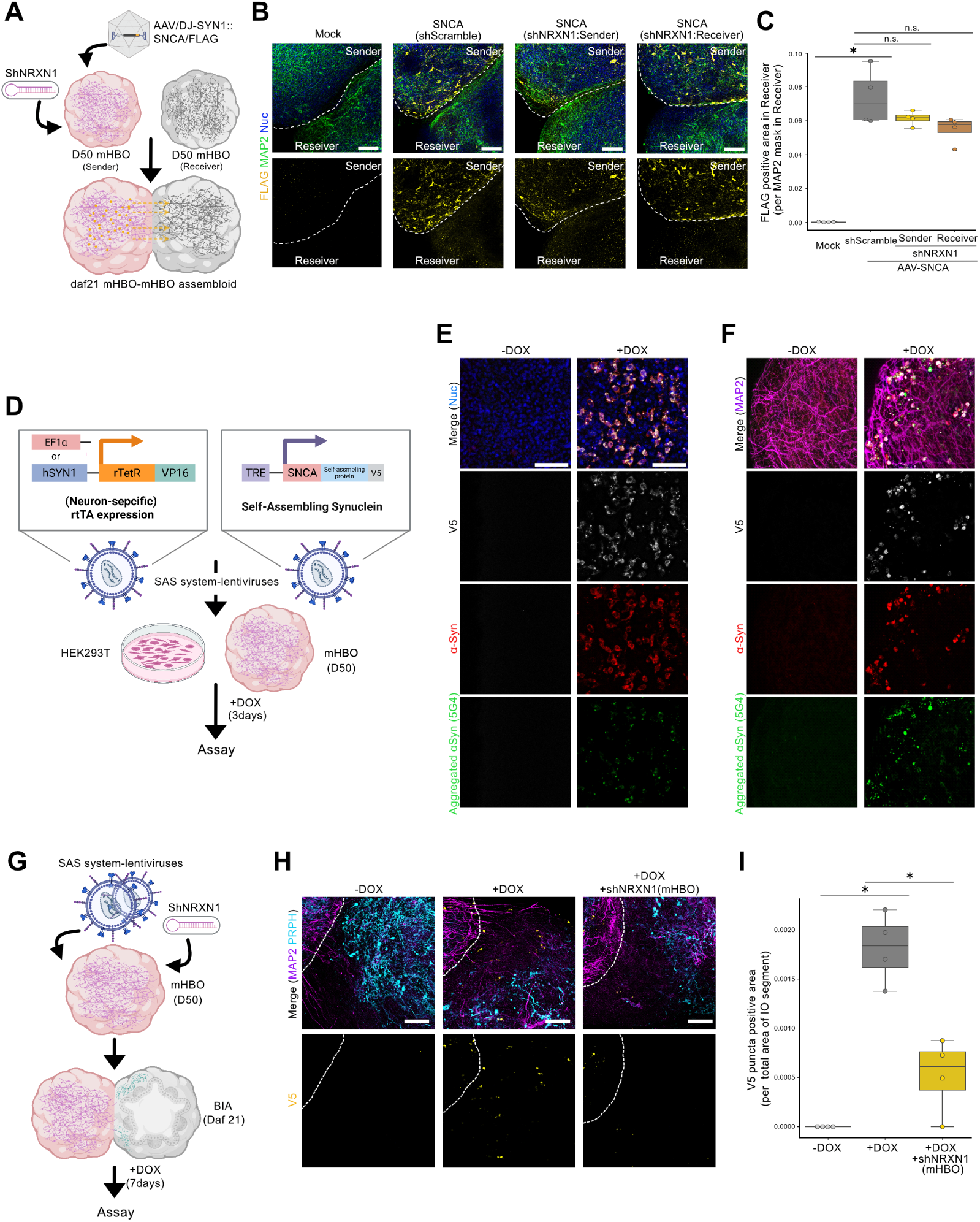
Additional validation of NRXN1 involvement using mHBO–mHBO assembloids and self-assembling α-syn, related to Figure 5. (A) Schematic of the generation of mHBO–mHBO assembloid and α-syn propagation assay. (B) Immunofluorescence staining of daf 21 mHBO–mHBO assembloids with or without shNRXN1. The dotted line indicates the boundary between sender and receiver regions. Scale bars, 100 μm. (C) Quantification of FLAG-positive area in receiver regions in (B). Data are shown as box-and-whisker plots (n = 4). Mann–Whitney test was used to compare Mock and AAV-SNCA (Mock), and Steel test was used for comparison among AAV-SNCA conditions. \**P* < 0.05. n.s., not significant. (D) Schematic of the self-assembling α-synuclein (SAS) system. (E) Immunofluorescence staining of SAS lentivirus–transduced HEK293T cells with or without doxycycline (DOX). Scale bars, 100 μm. (F) Immunofluorescence staining of SAS lentivirus–transduced mHBO with or without DOX. Scale bars, 100 μm. (G) Schematic of the SAS propagation assay in BIAs. (H) Immunofluorescence staining of SAS lentivirus-transduced BIAs with or without DOX and/or shNRXN1. Scale bars, 100 μm. (I) Quantification of V5-positive area in IO regions in (H). Data are shown as box-and-whisker plots (n = 4). The two-tailed Mann–Whitney test was used for comparison. \**P* < 0.05

**Figure S7.**
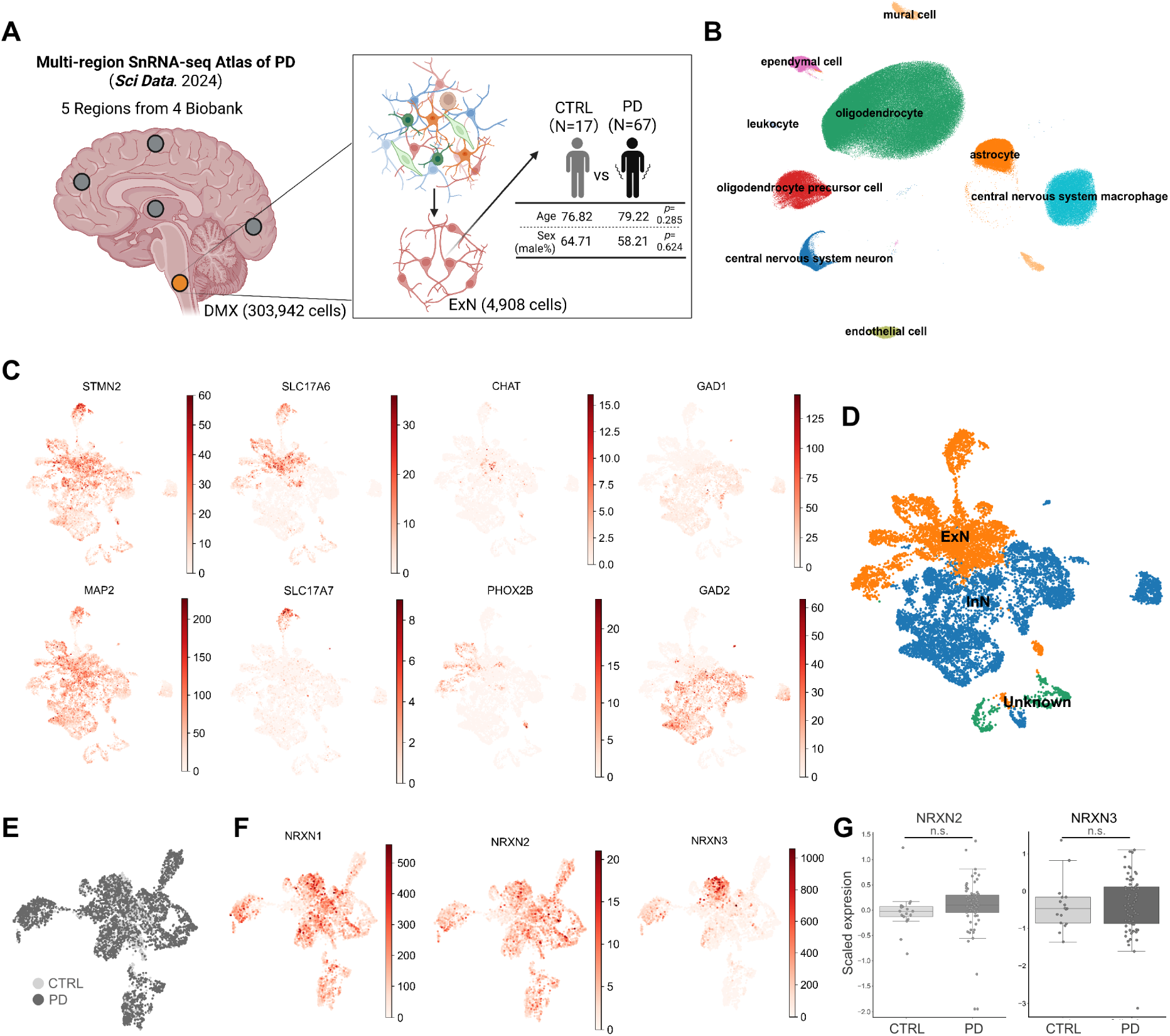
Reanalysis of multi-region single-nucleus RNA-seq atlas of PD. (A) Schematic of the reanalysis workflow for the multi-region single-nucleus RNA-seq atlas of PD. DMX-annotated cells were extracted for downstream analysis (left). After quality-filtering, the dataset comprised 4,908 cells from 17 healthy controls and 67 PD patients with comparable age and sex distributions (right). (B) UMAP embedding of the filtered dataset with cell-type annotations transferred from the original study. (C) Feature plots showing expression of representative neuronal subtype marker genes. (D) UMAP embedding with the identified neuronal subtypes. (E) UMAP embedding of ExN populations colored by condition. (F) Feature plots showing expression of NRXN1, NRXN2, and NRXN3. (G) NRXN2 and NRXN3 expression in ExN from control and PD samples. Data are shown as box-and-whisker plots (n = 17 controls, n = 67 PD). An unpaired two-tailed t-test was used for comparison. n.s., not significant.

## METHODS

KEY RESOURCES TABLE

| REAGENT OR RESOURCE | SOURCE | IDENTIFIER |
| --- | --- | --- |
| Antibodies |  |  |
| Goat Anti-OTX2 | R&D Systems | Cat#AF1979 |
| Mouse Anti-GBX2 | Sigma | Cat#SAB1403854 |
| Rabbit Anti-HOXB4 | Abcam | Cat#ab133521 |
| Mouse Anti-HOXC6 | Santa Cruz | Cat#sc-376330 |
| Mouse Anti-PHOX2B | Santa Cruz | Cat#sc-376997 |
| Goat Anti-CHAT | Sigma | Cat#AB144P |
| Mouse Anti-TUBB3 | Sigma | Cat#T8660 |
| Rabbit Anti-TUBB3 | BioLegend | Cat#802001 |
| Mouse Anti-VGLUT2 | Sigma | Cat#MAB5504 |
| Rabbit Anti-GABA | Sigma | Cat#A2052 |
| Rabbit Anti-SOX10 | Cell Signaling Technology | Cat#89356 |
| Mouse Anti-PRPH | Santa Cruz | Cat#sc-377093 |
| Rabbit Anti-PRPH | Cell Signaling Technology | Cat#85624 |
| Mouse Anti-SOX17 | Thermo Fisher Scientific | Cat#MA5-24885 |
| Goat Anti-FOXA2 | Santa Cruz | Cat#sc-6554 |
| Mouse Anti-SOX2 | Abcam | Cat#ab79351 |
| Rabbit Anti-CDX2 | Cell Signaling Technology | Cat#12306 |
| Rabbit Anti-CDH1 | R&D Systems | Cat#AF648 |
| Rabbit Anti-mCherry | Abcam | Cat#ab167453 |
| Rabbit Anti-pan-RFP | Proteintech | Cat#pabr1 |
| Chicken Anri-MAP2 | Abcam | Cat# 5392 |
| Mouse Anti-SNCA | Abcam | Cat# ab1903 |
| Rabbit Anti-SNCA | Abcam | Cat# ab138501 |
| Mouse Anti-FLAG | Sigma | Cat# F1804 |
| Mouse Anti-c-FOS | Abcam | Cat# ab208942 |
| Mouse Anti-V5 | Thermo Fisher Scientific | Cat# R960-25 |
| Mouse Anti-aggregated a-synuclein | Sigma | Cat# MABN389 |
| Chemicals, peptides, and recombinant proteins |  |  |
| AK02N | Ajinomoto | Cat# AK02N |
| iMatrix-511 | Matrixome | Cat# 892012 |
| TrypLE Select | Thermo Fisher Scientific | Cat# 12563011 |
| DMEM/F12 | Wako | Cat# 048-29785 |
| KnockOut Serum Replacement | Thermo Fisher Scientific | Cat# 10828-028 |
| Non-Essential Amino Acid | Sigma | Cat# M7145 |
| L-Glutamine | Nacalai | Cat# 16498-04 |
| β-Mercaptoethanol | Sigma | Cat# M3148 |
| Penicillin-Streptomycin | Nacalai | Cat# 6252-94 |
| GlutaMAX | Thermo Fisher Scientific | Cat# 35050061 |
| Dorsomorphin | Nacalai | Cat# 1876804 |
| SB431542 | Nacalai | Cat# 18176-54 |
| FGF2 | PeproTech | Cat# 100-18B |
| CHIR99021 | Nacalai | Cat# 18764-44 |
| Retinoic acid | Sigma | Cat# R2625 |
| Y27632 | Wako | Cat# 034-24024 |
| N2 Supplement | Wako | Cat# 141-09041 |
| KBM Neural Stem Cell | Cosmo Bio | Cat# 16050100 |
| B-27 Supplement minus vitamin A | Thermo Fisher Scientific | Cat# 12587010 |
| B-27 Plus Supplement | Thermo Fisher Scientific | Cat# A3582801 |
| BDNF | BioLegend | Cat# 788904 |
| GDNF | Alomone Labs | Cat# G-240 |
| L-Ascorbic Acid | Sigma | Cat# 5960 |
| Geltrex | Thermo Fisher Scientific | Cat# A1413202 |
| Amphotericin B | Sigma | Cat# A2942 |
| IWR1e | Sigma | Cat# 681669 |
| BMP4 | Qkine | Cat# QK038 |
| DAPT | Sigma | Cat# D5942 |
| RPMI1640 | Wako | Cat# 189-02145 |
| Activin A | Wako | Cat# 010-27623 |
| FGF-4 | Qkine | Cat# QK004 |
| EGF | PeproTech | Cat# AF-100-15 |
| R-spondin 1 | Qkine | Cat# QK006 |
| Paraformaldehyde | Nacalai | Cat# 26126-54 |
| CUBIC-L | TCI | Cat# T3740 |
| Triton X-100 | Wako | Cat# 169-21105 |
| Tween-20 | Wako | Cat# 163-21625 |
| TransIT-293 | Mirus | Cat# MIR2704 |
| Polybrene | VectorBuilder | Cat# PL200 |
| Polyethyleneimine | TCI | Cat# P0381 |
| BrainPhys Imaging Optimized Medium | STEMCELL Technologies | Cat# ST-05796 |
| Deschloroclozapine | Selleck | Cat# E1265 |
| IPTG | Wako | Cat# 096-05861 |
| Critical commercial assays |  |  |
| RNeasy mini kit | Qiagen | Cat# 74104 |
| SuperScript III First-Strand Synthesis SuperMix for qRT-PCR | Thermo Fisher Scientific | Cat# 11752050 |
| TB Green Premix Ex Taq (Tli RNaseH Plus) | Takara | Cat# RR420S |
| Papain Dissociation System | Worthington Biochemical | Cat# LK003150 |
| Chromium Next GEM Single Cell Fixed RNA Sample Preparation Kit | 10x Genomics | Cat# PN-1000414 |
| AAVpro Purification Kit Midi | Takara | Cat# 6675 |
| AAVpro Titration Kit (for Real Time PCR) Ver.2 | Takara | Cat# 6233 |
| Lenti-X Concentrator | Takara | Cat# 631231 |
| Lenti-X qRT-PCR Titration Kit | Takara | Cat# 631235 |
| Deposited data |  |  |
| Bulk RNA-seq | This paper | EBI: Will be listed upon publication |
| scRNA-seq | This paper | EBI: Will be listed upon publication |
| Bulk RNA-seq for Brainstem organoids | Eura et al., 2020<sup>24</sup> | DDBJ: DRA009864 |
| Bulk RNA-seq for mSpVO | Pang et al., 2024<sup>25</sup> | GEO: GSE251680 |
| Bulk RNA-seq for SCO | Chooi et al., 2022<sup>26</sup> | EBI: E-MTAB-12270 |
| Bulk RNA-seq for CO | Rosebrock et al., 2022<sup>27</sup> | GEO: GSE189981 |
| scRNA-seq for developing human brain | Braun et al., 2023<sup>29</sup> | https://github.com/linnarsson-lab/developing-human-brain/ |
| scRNA-seq for human gut cell atlas | Elmentaite et al., 2021<sup>50</sup> | https://cellxgene.cziscience.com/e/80a2c5b6-02e7-4fc0-9f12-179f5247c1bc.cxg/ |
| scRNA-seq for multi-region PD brains | Prashant et al., 2024<sup>53</sup> | https://cellxgene.cziscience.com/collections/d5d0df8f-4eee-49d8-a221-a288f50a1590 |
| Experimental models: Cell lines |  |  |
| hiPSC: 201B7 | Takahashi et al., 2007<sup>65</sup> | RRID: CVCL_A324 |
| hiPSC: WD39 | Imaizumi et al., 2012<sup>66</sup> | RRID: CVCL_Y528 |
| hiPSC: JB6 | Ishikawa et al., 2024<sup>67</sup> | RRID: CVCL_D0GW |
| Oligonucleotides |  |  |
| Primers for qRT-PCR, see Table S1. | This paper | N/A |
| Recombinant DNA |  |  |
| pAdDeltaF6 | N/A | Addgene plasmid# 11286 |
| AAV-DJ Rep-Cap | Cell Biolabs | Cat# VPK-420-DJ |
| pAAV-hSYN1-tdTomato | This paper | VectorBuilder# VB251216-1144xcu |
| pAAV-hSYN1-hM3D(Gq)-tagBFP2 | This paper | VectorBuilder# VB260101-1117ezk |
| pAAV-hSYN1-hSNCA/FLAG-tdTomato | This paper | VectorBuilder# VB240506-1269rhr |
| Lentiviral High Titer Packaging Mix | Takara | Cat# 6194 |
| pLV- EF1α-mCherry | This paper | VectorBuilder# VB010000-9492agg |
| pLV- hSYN1-jGCaMP8s | This paper | VectorBuilder# VB250605-1209bya |
| pLV- EF1α-rTetR | This paper | VectorBuilder# VB900167-2259ddw |
| pLV- hSYN1-rTetR | This paper | VectorBuilder# VB250707-1689rcx |
| pLV- TRE-SNCA/self-assembling protein/V5 | This paper | VectorBuilder# Will be listed upon publication |
| pLV- LacI-U6/2xLacO -shNRXN1 | This paper | VectorBuilder# VB250427-1086gry |
| pLV- LacI-U6/2xLacO -shNLGN2 | This paper | VectorBuilder# VB250922-1079qpt |
| pLV- LacI-U6/2xLacO -shScramble | This paper | VectorBuilder# VB010000-9535scz |
| Software and algorithms |  |  |
| ImageJ | National Institutes of Health | https://imagej.net/ |
| AGAVE | Allen Institute for Cell Science | https://github.com/allen-cell-animated/agave/releases/tag/v1.8.1 |
| Cutadapt (v1.9.1) | Martin et al., 2011<sup>68</sup> | https://github.com/marcelm/cutadapt |
| HISAT2 (v2.2.1) | Kim et al., 2019<sup>69</sup> | https://github.com/DaehwanKimLab/hisat2 |
| HTSeq (v0.6.1) | Anders et al., 2010<sup>70</sup> | https://github.com/htseq/htseq |
| sva (v3.50.0) | Johnson et al., 2007<sup>71</sup> | https://bioconductor.org/packages/release/bioc/html/sva.html |
| DESeq2 (v1.42.1) | Love et al., 2014<sup>72</sup> | https://bioconductor.org/packages/release/bioc/html/DESeq2.html |
| GSVA (v 1.50.5) | Hänzelmann et al., 2013<sup>73</sup> | https://github.com/rcastelo/GSVA |
| Voxhunt (v 1.0.1) | Fleck et al., 2020<sup>74</sup> | https://github.com/quadbio/VoxHunt |
| ggplot2 (v3.5.0) | N/A | https://ggplot2.tidyverse.org/ |
| Cell Ranger (v 7.0.1) | N/A | https://www.10xgenomics.com/support/software/cell-ranger |
| Seurat (v 4.3.0) | Hao et al., 2021<sup>75</sup> | https://satijalab.org/seurat/ |
| Harmony (v 1.2.3) | Korsunsky et al., 2019<sup>76</sup> | https://github.com/immunogenomics/harmony |
| AUCell (v 1.24.0) | Aibar et al., 2017<sup>77</sup> | https://bioconductor.org/packages/release/bioc/html/AUCell.html |
| URD | Farrel et al., 2018<sup>78</sup> | https://github.com/farrellja/URD |
| CellChat (v2) | Jin et al., 2025<sup>45</sup> | https://github.com/SiYangming/CellChat2 |
| Scanpy (v1.11.5) | Wolf et al., 2018<sup>79</sup> | https://scanpy.readthedocs.io/en/stable/index.html |
| Seaborn (v 0.13.2) | N/A | https://seaborn.pydata.org/ |
| JMP Pro (v18) | JMP | N/A |
| Other |  |  |
| Ultra-low attachment 96-well plate | Sumitomo Bakelite | Cat# MS-9096M |
| Ultra-low attachment 6-well plate | Corning | Cat# 3471 |
| 35-mm glass-bottom dish | Matsunami Glass | Cat# D11131H |
| BZ-X810 microscope | Keyence | Cat# BZ-X810 |
| CellVoyager CQ1 | Yokogawa | N/A |
| QuantStudio 7 Flex Real-Time PCR System | Thermo Fisher Scientific | Cat# QS7-03 |

## METHOD DETAILS

### Cell lines

Healthy control human iPSC lines 201B7^65^, WD39^66^, and JB6^67^ were cultured on iMatrix-511-coated tissue culture dishes with AK02N media in a humidified 37 °C incubator with 5% CO_2_. Human pluripotent stem cells (hPSCs) were passaged every 7 days by 0.5x TrypLE Select diluted in PBS(−). All experimental procedures involving human iPSCs were approved by the Juntendo University School of Medicine Ethics Committee (approval no. 2017032).

### Generation of mHBOs

On day 0, feeder-free hiPSC colonies were dissociated into single cells using 0.5x TrypLE Select treatment for 5 min at 37 °C. Cells were resuspended in KSR induction media consisting of DMEM/F12, supplemented with 20% (v/v) KSR, 1% (v/v) MEM-NEAA, 2mM L-Glutamine, and 100 μM β-Mercaptoethanol and 1% (v/v) penicillin–streptomycin. The medium was further supplemented with 5µM Dorsomorphin, 10 μM SB431542, 20 ng/mL FGF2, and 10 μM Y27632 were plated to an ultra-low attachment 96-well plate (2×10^4^ cells/well) centrifuged at 200 *g* for 1 min, and then incubated at 37 °C in 5% CO_2_.

On day 6, the media was replaced to N2 medium composed of DMEM-F12, supplemented with 1% (v/v) N2, 0.5% (v/v) MEM-NEAA, 1% (v/v) GlutaMAX, and 1% (v/v) penicillin–streptomycin, together with 5µM Dorsomorphin, 10 μM SB431542, and 20 ng/mL FGF2. For caudalization and hindbrain patterning, 1–3 μM CHIR99021 and/or 100–3000 nM retinoic acid was added to the induction medium during days 0–11, as indicated for each experimental condition.

On day 11, the medium was changed to differentiation medium consisting of KBM Neural stem cell 2% (v/v) B-27 supplement minus vitamin A 1% (v/v) Geltrex and 1% (v/v) penicillin–streptomycin, together with 20 ng/mL BDNF, 20 ng/mL GDNF and 200 μM ascorbic acid. The medium was replenished every 5 days, and Geltrex was omitted from the medium after day 21.

From day 30 onward, organoids were maintained in maturation medium consisting of KBM Neural Stem Cell medium supplemented with 2% (v/v) B-27 Plus supplement, and 1% (v/v) penicillin–streptomycin, 20 ng/mL BDNF, 20 ng/mL GDNF, 200 μM ascorbic acid, and 0.5 μg/mL Amphotericin B. The medium was replenished every 5 days thereafter.

mHBO–mHBO assembloids were generated by fusing day 50 mHBOs. Individual organoids were placed in the same well of an ultra–low–attachment 96–well plate and centrifuged at 800 × g for 2 min to promote fusion. The assembloids were maintained in maturation medium. The medium was replenished every four days thereafter.

### Generation of non-patterned brain organoids and cortical organoids

On day 0, feeder-free hiPSC colonies were dissociated into single cells using 0.5× TrypLE Select treatment for 5 min at 37 °C. Cells were resuspended in KSR induction medium supplemented with 5 µM Dorsomorphin, 10 μM SB431542, 10 μM Y27632, and 3µM IWR1e (cortical organoids only), and plated into an ultra-low attachment 96-well plates (4×10^4^ cells/well) centrifuged at 200 × *g* for 1 min.

On day 6, the medium was replaced with N2 medium supplemented with 5 µM Dorsomorphin, 10 μM SB431542, and 3µM iwr1e (cortical organoids only).

On day 11, the medium was changed to differentiation medium supplemented with 20 ng/mL BDNF, 20 ng/mL GDNF, and 200 μM ascorbic acid. The medium was replenished every 5 days thereafter.

### Generation of ENCCs

ENCCs were generated as previously described,^39^ with minor modifications. Briefly, hiPSC colonies were dissociated into single cells using 0.5× TrypLE Select for 5 min at 37 °C and plated onto 1% Geltrex-coated 6-well plates at a density of 3 × 10⁵ cells per well. Cells were cultured in N2 induction medium supplemented with 1 μM dorsomorphin, 2 μM SB431542, and 20 ng/mL BMP4. Y-27632 (10 μM) was added only on day 0.

On day 1, the medium was replaced with fresh induction medium. On day 4, the medium was replaced with induction medium supplemented with 1 μM retinoic acid, and the same medium was refreshed again on day 5.

On day 7, cells were dissociated using 0.5× TrypLE Select and transferred to suspension culture in an ultra-low attachment 6-well plate to form spheroids in spheroid medium (KBM Neural Stem Cell medium supplemented with 2% (v/v) B-27 minus vitamin A and 1% (v/v) penicillin–streptomycin) supplemented with 3 µM CHIR99021, 20 ng/mL FGF2, and 10 µM Y27632.

To assess differentiation efficiency, day 11 spheroids were collected into 1.5-mL microcentrifuge tubes and allowed to settle by gravity for 5 min. After removal of the supernatant, spheroids were resuspended in differentiation medium (KBM Neural Stem Cell medium supplemented with 2% (v/v) B-27 minus vitamin A 1% (v/v) penicillin–streptomycin) supplemented with 20 ng/mL BDNF, 20 ng/mL GDNF, 200 μM ascorbic acid, and 10 μM DAPT. The medium was replenished every 3 days thereafter.

### Generation of IOs

IOs were generated as previously described,^40^ with minor modifications. Briefly, hiPSC colonies were dissociated into single cells using 0.5× TrypLE Select for 5 min at 37 °C and plated onto iMarix-511–coated 12-well plates at a density of 1 × 10^5^ cells per well (day-7). Cells were maintained in AK02N medium until differentiation was initiated.

On day 0, the medium was replaced with RPMI 1640 supplemented with 100 ng/mL Activin A and 3 μM CHIR99021. On day 1, the medium was changed to RPMI 1640 supplemented with 100 ng/mL Activin A and 1% (v/v) KSR. On day 2, the medium was replaced with RPMI 1640 supplemented with 100 ng/mL Activin A and 10% (v/v) KSR.

From day 4 onward, the medium was changed every other day to RPMI 1640 supplemented with 2 mM GlutaMAX, 10% (v/v) KSR, 3 μM CHIR99021, and 100 ng/mL FGF4.

On day 11, cells were dissociated using 0.5× TrypLE Select and resuspended in intestinal organoid medium (DMEM/F12 supplemented with 2 mM GlutaMAX, 2% (v/v) B-27 minus vitamin A, and 1% (v/v) penicillin–streptomycin) supplemented with 100ng/mL EGF, 500 ng/mL R-Spondin1, and 10µM Y27632. Cells were plated into ultra-low-attachment 96-well plates (5×10^4^ cells/well), centrifuged at 200 × g for 1 min, and incubated at 37 °C in 5% CO₂.

On day 14, the medium was replaced with intestinal organoid medium supplemented with 100 ng/mL EGF, and the same medium was refreshed again on day 16. From day 18 onward, organoids were maintained in intestinal organoid maturation medium (DMEM/F12 supplemented with 2 mM GlutaMAX, 2% (v/v) B-27 Plus supplement. and 1% (v/v) penicillin–streptomycin) supplemented with 100ng/mL EGF. The medium was replenished every 4 days thereafter.

### Generation of IOs with enteric neurons

To generate IOs containing enteric neurons, day 11 ENCC spheroids were collected and dissociated using 0.5× TrypLE Select for 5 min at 37 °C. In selected experiments, ENCCs were labeled at day 10 by lentiviral transduction with an EF1α-driven mCherry reporter prior to dissociation. Dissociated ENCCs were added at a density of 2 × 10^5^ cells per well to day 9 hindgut cultures and co-cultured in RPMI 1640 supplemented with 2 mM GlutaMAX, 10% (v/v) KSR, 3 μM CHIR99021, and 100 ng/mL FGF4.

On day 11, hindgut cultures containing ENCCs were dissociated and plated into ultra-low-attachment 96-well plates to generate enteric neuron–enriched IOs, following the same spheroid formation and maturation protocol used for intestinal organoids, except that the intestinal organoid medium was additionally supplemented with 20 ng/mL GDNF and 200 μM ascorbic acid.

### Generation of BIAs

To generate BIAs, day 50 mHBOs were fused with day 21 IOs, with or without ENCC incorporation. Individual organoids were placed together in ultra-low-attachment 96-well plates and centrifuged at 800 × g for 2 min. Assembloids were maintained in an assembloid medium (DMEM/F12 supplemented with 2 mM GlutaMAX, 2% (v/v) B-27 Plus supplement, 1% (v/v) penicillin–streptomycin, and 0.5 μg/mL Amphotericin B) supplemented with 100 ng/mL EGF, 20 ng/mL GDNF, and 200 μM ascorbic acid. The medium was replenished every four days thereafter.

### Whole-mount immunofluorescence staining

Organoids and assembloids were fixed in 4% paraformaldehyde (PFA) at 24 °C for 3 h and washed three times with PBS (−). Organoids and assembloids were fixed in 4% PFA at 24℃ for 3 h, washed 3 times with PBS(-). Delipidation was performed using CUBIC-L only for samples shown in Figure 3M. Blocking and permeabilization were carried out in PBS containing 10% fetal bovine serum (FBS) and 0.3% Triton X-100 at 24 °C for 3 h.

Samples were then incubated with primary antibodies diluted in PBS containing 10% FBS and 0.3% Triton X-100 for two nights at 4 °C. After three washes with 0.05% Tween-20 in PBS, samples were incubated with appropriate secondary antibodies diluted in the same buffer for an additional overnight incubation at 4 °C. Samples were subsequently washed three times with 0.05% Tween-20 in PBS and transferred onto glass slides.

For tissue clearing, FunGI^80^ was applied at 24 ℃ for 1 h. A list of antibodies is provided in the Key Resources Table. Fluorescence images were acquired using a Keyence BZ-X810 microscope and analyzed with ImageJ. Three-dimensional (3D) rendering was performed using AGAVE.

### Real-time quantitative PCR (qPCR)

For quantitative PCR analysis, 12–24 organoids were pooled per sample, and total RNA was isolated using the RNeasy Mini Kit with QIAshredder homogenization, according to the manufacturer’s instructions. cDNA was synthesized using the SuperScript III First-Strand Synthesis SuperMix for qRT-PCR. qPCR was performed using TB Green Premix Ex Taq (Tli RNaseH Plus) on a QuantStudio 7 Flex Real-Time PCR System.

The PCR cycling conditions were as follows: 95 °C for 30 s, followed by 40 two-step cycles at 95 °C for 5 s, and 60 °C for 30 s. Values were normalized to ACTB and analyzed using the comparative (ΔΔCt) method. Primer sequences used in this study are provided in Table S1.

### Bulk RNA-seq and data analysis

Day 11–100 organoids (16–24 organoids per sample) were collected, and total RNA was isolated using the RNeasy Mini Kit with QIAshredder homogenization, according to the manufacturer’s instructions. cDNA libraries were constructed using the TruSeq stranded mRNA Library Preparation kit and sequenced on a NovaSeq6000 platform to obtain 150 bp paired-end reads (Azenta).

Raw sequencing reads were processed for quality control using Cutadapt (v1.9.1) to remove adapter sequences and low-quality bases (Phred score < 20). Clean reads were aligned to the human reference genome (GRCh38) using HISAT2 (v2.2.1). Gene-level expression quantification was performed using HTSeq (v0.6.1) based on annotated gene models.

Published RNA-seq data from organoid samples were downloaded from the NCBI Sequence Read Archive (SRA), the DNA Data Bank of Japan (DDBJ), and the European Nucleotide Archive (ENA). For integrative analyses across datasets, batch effects were corrected using ComBat-seq implemented in the sva package (v3.50.0). Normalized gene expression values were converted to transcripts per kilobase of exon model per million mapped reads (TPM). Heatmaps of gene expression were generated using row-wise z-scores of log_2_(TPM +1) for each selected gene.

Differential gene expression analysis was performed using DESeq2 (v1.42.1), and statistical significance was assessed using Wald tests. P–values were adjusted for multiple testing using the Benjamini–Hochberg method. Transcriptomic similarity analysis was performed using the single-sample gene set enrichment analysis (ssGSEA) implemented in the GSVA package (v 1.50.5). Brain region–specific gene signatures were obtained from the top 20 genes calculated by Voxhunt (v 1.0.1), as described previously.^25^ Enrichment scores were min–max normalized and visualized as heatmaps using ggplot2 (v3.5.0).

### scRNA-seq sample preparation and library construction

Day 70 organoids or daf 20 assembloids (20–30 organoids per sample) were dissociated into single-cell suspensions using the Papain Dissociation System. Samples were incubated in 20 U/mL papain enzyme solution supplemented with 10,000 U/mL DNase I at 37 °C for 45 min, with gentle trituration every 10 min to facilitate dissociation. Following enzymatic digestion, dissociation was stopped by washing the samples with a protease inhibitor–containing solution, and cells were gently triturated to obtain a single-cell suspension. The suspension was then filtered through a 70 µm cell strainer to remove debris and undissociated aggregates, and cell numbers were determined.

Single-cell suspensions were fixed using the Chromium Next GEM Single Cell Fixed RNA Sample Preparation Kit, according to the manufacturer’s instructions. Fixed cells were resuspended in Quenching Buffer supplemented with Enhancer and 10% glycerol and stored at −80 °C until downstream processing. Fixed single-cell suspensions were processed using the Chromium Next GEM Single Cell Fixed RNA Library v1 – Human 4BC kit, according to the manufacturer’s instructions. The libraries were sequenced on a NovaSeq platform using 150-bp paired-end reads (Macrogen).

### Preprocessing, quality control, and cell-type annotation

Sequencing reads were aligned to the human genome GRCh38 (v 3.0.0, provided by 10x genomics) using Cell Ranger (v 7.0.1) with default parameters. Downstream analysis was performed using Seurat (v 4.3.0). For quality control, cells with 500–10000 detected genes, 2000–50000 unique molecular identifiers (UMIs), and those with <10% mitochondrial gene fraction were retained as high-quality cells.

Gene expression was normalized using default parameters, and the 2,000 most variable genes were selected (selection.method = “vst”) and scaled prior to principal component analysis (PCA). The top 20 principal components (PCs) were used for clustering (resolution = 1.0) for visualization with UMAP.

Mesenchymal populations, defined by COL1A1 and DCN expression, were excluded prior to cell–type annotation, specifically in the hindbrain organoid datasets. Clusters were annotated based on the expression of established marker genes, as described previously.^28^

For assembloid datasets, cell-type annotations were transferred from the hindbrain organoid dataset, whereas intestinal organoid–derived populations were annotated based on marker gene expression. For integrative analyses across datasets, batch effects were corrected using Harmony (v1.2.3).

To compare organoid-derived cells with published developing human brain atlas scRNA-seq datasets^29^, anchor-based label transfer was performed using Seurat. Briefly, 100,000 cells were randomly subsampled from the combined datasets, anchors between the query and reference datasets were identified and filtered, and predicted cell-type labels were projected onto the UMAP embedding.

To compute spatial correlation patterns between the single-cell transcriptome and medullary dorsoventral subregions, the AUCell package (v 1.24.0) was used to calculate enrichment scores for each cell type, which were subsequently Z-scaled for visualization.

### Pseudotime analysis and lineage trajectory analysis using URD

Pseudotime analysis was performed using URD.^78^ Gene expression matrices, highly variable genes, PCA loadings, and UMAP embeddings were imported from the Seurat object into a URD object. Diffusion maps were computed using principal component space (k = 90), and radial glial cells were defined as the root node.

Pseudotime values were assigned using the flood-based pseudotime algorithm implemented in URD, and the robustness of pseudotime ordering was assessed using built-in stability metrics. Terminal cell populations were manually defined based on cluster identity. A pseudotime-biased transition matrix was generated using logistic regression–based weighting to enforce directionality along developmental time.

Developmental trajectories were reconstructed by biasing transition probabilities along pseudotime and simulating random walks from terminal tip clusters to the root. A branching tree was built based on visitation frequencies and divergence preference.

### Ligand–receptor analysis

Cell–cell communication analysis was performed using CellChat v2^45^ with the human ligand–receptor interaction database (CellChatDB.human). Overexpressed genes and ligand–receptor interactions were identified, and communication probabilities were computed using the default CellChat workflow. Interactions supported by fewer than 10 cells were filtered out.

Communication probabilities were summarized at both ligand–receptor pair and signaling pathway levels, and global communication networks were constructed. For comparative analyses between conditions, CellChat objects were merged, and differential interaction strength and pathway activity were quantified using built-in comparison functions. Changes in signaling strength, pathway usage, and cell-type–specific signaling roles were visualized using differential network plots, heatmaps, and ranked signaling analyses.

### Reanalysis of publicly available datasets

The human gut cell atlas dataset was downloaded from Cellxgene and reanalyzed using Scanpy (v1.11.5). Cells annotated as adult samples in the original metadata were selected for downstream analysis. Gene expression was normalized using the default parameters, and the 2,000 most variable genes were selected and scaled prior to PCA. The top 50 PCs were used for clustering (resolution = 0.1) and visualization with UMAP. Cell-type annotations were transferred from the original dataset, and gene expression patterns were visualized using UMAP and dot plots.

A publicly available multi-region single-nucleus RNA-seq atlas of Parkinson’s disease brains was also downloaded from Cellxgene and analyzed using Scanpy. Nuclei derived from the dorsal motor nucleus of the vagus nerve in the original metadata were extracted. ’Glutamatergic neuron’ and ’GABAergic neuron’ were excluded owing to their limited representation in the dataset. All remaining annotated cell types were initially projected onto the UMAP embedding, after which nuclei annotated as central nervous system neurons were retained for further analysis. Gene expression was normalized using default parameters, and the 2,000 most variable genes were selected and scaled prior to PCA. The top 30 PCs were used for clustering (resolution = 0.2) and visualization with UMAP. Clusters were annotated based on marker gene expression. Among neuronal subtypes, excitatory neuronal populations were extracted, and the remaining neuronal populations were reprocessed, including normalization, variable gene selection, PCA, and clustering (resolution = 0.1). Gene expression profiles were compared between healthy control and PD samples and visualized using UMAP and box plots.

### Viral labeling

For AAV production, HEK293T cells were co-transfected using TransIT-293 Reagent according to the manufacturer’s instructions with pAdDeltaF6, AAV-DJ Rep-Cap plasmid, and the respective vector plasmids (hSYN1-tdTomato, hSYN1-hM3D(Gq)-tagBFP2, hSYN1-hSNCA/FLAG-tdTomato). AAV particles were purified using the AAVpro® Purification Kit Midi, and viral titers were determined by quantitative PCR using the AAVpro® Titration Kit (Real-Time PCR) Ver.2, following the manufacturer’s protocols.

For lentivirus production, HEK293T cells were co-transfected using TransIT-293 Reagent with the Lentiviral High Titer Packaging Mix and the corresponding vector plasmids (EF1α-mCherry, hSYN1-jGCaMP8s, EF1α-rTetR, hSYN1-rTetR, TRE-SNCA/self-assembling protein/V5, or LacI-U6/2xLacO - shNRXN1/shNLGN2/shScramble). Self-assembling α-synuclein (SAS) vectors were generated based on amino acid sequences reported by Fan et al.^52^ and codon-optimized prior to cloning. Lentiviral particles were concentrated from culture supernatants using Lenti-X Concentrator, and viral titers were quantified using the Lenti-X qRT-PCR Titration Kit. For viral transduction, organoids were incubated with either AAV (4–8 × 10^8^ genome copies per organoid) or lentivirus (0.5–2 × 10^5^ transducing units per organoid) in culture medium. Polybrene (2.5 µg/mL) was added only for lentiviral transduction. Infections were performed in 96-well plates with a total volume of 50 µL per well, and organoids were incubated for two nights at 37 °C with 5% CO₂. Following transduction, organoids were washed twice with DMEM/F12 and standard culture conditions.

### Calcium imaging

For chemogenetic stimulation experiments, AAV/DJ-hSYN1-hM3D(Gq) was introduced into day 50 mHBOs prior to assembloid formation. Daf 28 assembloids were attached to 35-mm glass-bottom dishes coated with 0.1% polyethyleneimine (PEI) for live imaging. Assembloids were maintained in BrainPhys Imaging Optimized Medium supplemented with 2% (v/v) B-27 Plus supplement, 1% (v/v) penicillin–streptomycin, 20 ng/mL GDNF, and 200 µM ascorbic acid. For hM3D stimulation, 10 nM DCZ was applied to selective activate hM3D(Gq)-expressing neurons. Calcium dynamics were recorded using a CellVoyager CQ1 high-content imaging system. Calcium activity was recorded for 5 min prior to DCZ application at a frame rate of 5 frames per second. Regions of interest (ROI) were identified using semi-automated methods, and fluorescence intensities were quantified over time using CellPathfinder. To correct for gradual baseline decay, fluorescence traces were linearly detrended by estimating the slope between the minimum intensity values in the initial and final segments of each recording and subtracting this drift from the raw signal. Relative changes in intracellular calcium were calculated as ΔF/F₀, where F₀ was defined as the 5th percentile of the baseline-corrected fluorescence trace for each ROI. For the c–Fos expression assay, AAV/DJ–hSYN1–hM3D(Gq)–tagBFP2 was used, followed by the chemogenetic stimulation described above and subsequent fixation for immunostaining.

### α-syn Propagation assay

To assess brain-to-gut α-syn propagation, either AAV/DJ-hSYN1-hSNCA/FLAG-tdTomato or a lentiviral Tet-inducible self-assembling α-syn (SAS) system (hSYN1-rTetR and TRE-SNCA/self-assembling protein/V5) was introduced into day 50 mHBOs prior to assembloid formation. Virus-infected mHBOs were subsequently fused with IOs to generate BIAs. For SAS propagation experiments, assembloids were treated with 2 μg/mL doxycycline starting at day 14 after fusion and maintained until fixation. Daf 21 assembloids were fixed and processed for immunohistochemical analysis. Exogenous α-syn propagation was evaluated by detecting FLAG/V5-tagged α-syn signals in enteric neuronal filaments within the gut compartment.

### Knockdown experiments

For NRXN1 and NLGN2 knockdown, inducible shRNA–expressing lentiviral vectors were introduced into day 48 mHBOs, day 19 IOs, and/or day 10 ENCCs prior to assembloid formation. Virus–infected organoids were subsequently used to generate BIAs. For knockdown efficiency analysis, mHBOs and IOs were maintained in maturation medium containing 200 µM IPTG, whereas ENCCs were plated and cultured in differentiation medium containing 200 µM IPTG. In BIAs, IPTG treatment was initiated at day 7 after fusion (daf 7) and maintained until sample collection. Assembloids were collected at daf 21 and processed for immunostaining.

## QUANTIFICATION AND STATISTICAL ANALYSIS

Data are presented as box-and-whisker plots with individual data points overlaid. Statistical analyses were performed using JMP Pro (v18). Comparisons were conducted using two-tailed unpaired Student’s t-test, two-tailed Mann–Whitney tests, or Steel’s tests, as appropriate for each dataset. Differential gene expression analysis of RNA-seq data was performed using Wald tests. A *P* value < 0.05 was considered statistically significant. Data visualization was performed using seaborn (0.13.2).

