## Supplementary material for "Modeling the pathological brain-gut axis in Parkinson’s disease using human iPSC derived brain-intestinal assembloids": Table S1

**Table S1. Primers for RT-qPCR**

| Gene Symbol |  | Sequence (5' to 3') |
| --- | --- | --- |
| ACTB | Forward | TGAAGTGTGACGTGGACATC |
|  | Reverse | GGAGGAGCAATGATCTTGAT |
| EN1 | Forward | TGGGTGTACTGCACACGTTATTC |
|  | Reverse | GGAACTCCGCCTTGAGTCTCT |
| GBX2 | Forward | AAAGAGGGCTCGCTGCTC |
|  | Reverse | ATCGCTCTCCAGCGAGAA |
| HOXA2 | Forward | CGTCGCTCGCTGAGTGCCTG |
|  | Reverse | TGTCGAGTGTGAAAGCGTCGAGG |
| HOXB3 | Forward | AACGCCTTACACTCCATGACC |
|  | Reverse | ATTCTGGTGGGCTTTACCGAA |
| HOXB4 | Forward | CGTGAGCACGGTAAACCCCAA |
|  | Reverse | ATTCCTTCTCCAGCTCCAAGACCT |
| HOXC6 | Forward | TGAATTCCTACTTCACTAACCCTTC |
|  | Reverse | ATCATAGGCGGTGGAATTGA |
| PHOX2B | Forward | CTACCCCGACATCTACACTCG |
|  | Reverse | TCCTGCTTGCGAAACTTG |
| NRXN1 | Forward | TAAGTGGCCTCCTAATGACCG |
|  | Reverse | TCGCACCAATACGGCTTCTTT |
| NLGN2 | Forward | TCAACTACCGTCTTGGGGTG |
|  | Reverse | GTGGGCGATGTTTTCACTGAG |
